# Broad variation in response of individual introns to splicing inhibitors in a humanized yeast strain

**DOI:** 10.1101/2023.10.05.560965

**Authors:** Oarteze Hunter, Jason Talkish, Jen Quick-Cleveland, Haller Igel, Asako Tan, Scott Kuersten, Sol Katzman, John Paul Donohue, Melissa Jurica, Manuel Ares

## Abstract

Intron branch point (BP) recognition by the U2 snRNP is a critical step of splicing, vulnerable to recurrent cancer mutations and bacterial natural product inhibitors. The BP binds a conserved pocket in the SF3B1 (human) or Hsh155 (yeast) U2 snRNP protein. Amino acids that line this pocket affect binding of splicing inhibitors like Pladienolide-B (Plad-B), such that organisms differ in their sensitivity.

To study the mechanism of splicing inhibitor action in a simplified system, we modified the naturally Plad-B resistant yeast *Saccharomyces cerevisiae* by changing 14 amino acids in the Hsh155 BP pocket to those from human. This humanized yeast grows normally, and splicing is largely unaffected by the mutation. Splicing is inhibited within minutes after addition of Plad-B, and different introns appear inhibited to different extents. Intron-specific inhibition differences are also observed during co-transcriptional splicing in Plad-B using single-molecule intron tracking (SMIT) to minimize gene-specific transcription and decay rates that cloud estimates of inhibition by standard RNA-seq. Comparison of Plad-B intron sensitivities to those of the structurally distinct inhibitor Thailanstatin-A reveals intron-specific differences in sensitivity to different compounds. This work exposes a complex relationship between binding of different members of this class of inhibitors to the spliceosome and intron-specific rates of BP recognition and catalysis. Introns with variant BP sequences seem particularly sensitive, echoing observations from mammalian cells, where monitoring individual introns is complicated by multi-intron gene architecture and alternative splicing. The compact yeast system may hasten characterization of splicing inhibitors, accelerating improvements in selectivity and therapeutic efficacy.

## INTRODUCTION

Eukaryotic gene expression relies on accurate and timely splicing of pre-mRNA transcripts by the spliceosome. The spliceosome is an RNA-protein complex that assembles anew on each intron and proceeds through a litany of compositional and conformational changes as it executes the reactions necessary to remove the intron and ligate the flanking exons (for review, see Wilkinson et al. 2020). For the most part, the early assembly steps presage the splicing outcome for the mRNA. Recognition of the 5’ splice site (5’ss) by the U1 snRNP and the branch point (BP) by the U2 snRNP capture the intron reactive groups that will be gathered at the catalytic site by subsequent rearrangements. Selection of the 3’ splice site (3’ss) is dictated in large part by BP choice, and is very often found at the first YAG that is >7 nucleotides from the BP (Smith et al. 1993; Chua and Reed 2001). Since the early steps in spliceosome assembly have an outsized influence on the spliced product, much effort has been focused on understanding their mechanism and regulation.

The molecular events leading to stable association of the U2 snRNP with the intron BP are only partially understood. The E-complex, containing pre-mRNA and U1 snRNP (Michaud and Reed 1993, 1991; Li et al. 2019), must merge with a form of the U2 snRNP carrying the SF3a and SF3b proteins (called 17S U2 snRNP in human, Brosi et al. 1993b, 1993a) to recognize the BP and form the prespliceosome or A-complex, within which the 5’ss and BP are base paired to snRNAs U1 and U2, respectively (Zhang et al. 2020b; Plaschka et al. 2018). As the prespliceosome forms, proteins present in E-complex and the U2 snRNP are released to allow new RNA-RNA interactions to be established. For example, in the yeast E-complex, the intron BP region is bound by a heterodimer of Msl5/Bbp and Mud2 (Berglund et al. 1998; Li et al. 2019), and is inaccessible to its future RNA pairing partner, the BP interacting stem loop (BSL) of U2 snRNA (Perriman and Ares 2010). In yeast, genetic interactions between the U2AF65 homolog Mud2 and the RNA helicase family member Sub2 (UAP56, BAT1, or DDX39B in humans) suggests that the intron branch point might be cleared of proteins by Sub2 (Kistler and Guthrie 2001).

In the U2 snRNP, a U2 snRNA stem loop structure called the BSL (Perriman and Ares 2010) is bound by Cus2 (yeast), or TATSF1 (humans, Zhang et al. 2020b; Tholen et al. 2022), which must be removed to allow the BSL to open and pair with the intron to establish the U2-BP helix (Perriman and Ares 2010). In yeast, Cus2 is released by the ATP-dependent activity of the RNA helicase family member Prp5 (DDX46 in humans, Perriman et al. 2003; Talkish et al. 2019a). Once formed, the U2-BP helix is bound through conformational changes of Hsh155 (in humans, SF3B1), and the BP-adenosine (BP-A) residue enters the BP-A binding pocket formed by HEAT repeats 15 and 16 of this conserved U2 protein (Rauhut et al. 2016; Yan et al. 2016; Plaschka et al. 2018). In yeast, Prp5 releases from correctly formed U2-BP helices allowing splicing to proceed, but remains associated with malformed U2-BP helices, serving as a fidelity check (Xu and Query 2007) that prevents the tri-snRNP from binding (Liang and Cheng 2015; Kao et al. 2021). Once bound in the BP-A binding pocket of SF3B1/Hsh155, the BP is held awaiting arrival of the tri-snRNP to form the pre-B-complex. Upon activation of the B-complex by the Prp2 RNA helicase (DHX16 in humans), the SF3B proteins are destabilized, allowing the BP to dock in the active site for 5’ splice site cleavage and branch formation (Bai et al. 2021; Wan et al. 2019).

Correct BP recognition is critical for correct 3’ splice site selection, which is in turn essential for producing mRNA with the correct reading frame. The importance of correct BP selection is underscored by two additional findings. One is that BP selection is altered in cancerous cells carrying recurrent mutations in SF3B1 that influence tumor progression (Darman et al. 2015; Alsafadi et al. 2016), and the other is that bacteria produce small molecule splicing inhibitors (Spliceostatin-A, Pladienolide-B, etc.) that block loading of the branch point in to SF3B1 (Yokoi et al. 2011; Cretu et al. 2021). *In vitro*, formation of the prespliceosome or A-complex is blocked by this group of inhibitors (Corrionero et al. 2011; Roybal and Jurica 2010; Folco et al. 2011) as a consequence of their binding the BP-A binding pocket between HEAT repeats 15-16 of SF3B1 and PFH5A (Teng et al. 2017; Cretu et al. 2018, 2021; Finci et al. 2018). The relationships between the anti-tumor activity of splicing inhibitors and the contribution of recurrent mutations along a pathway of tumor evolution are difficult to attribute to changes in splicing of specific introns, however for many cancers, a general inhibition of splicing is selectively lethal, opening therapeutic opportunities (Hsu et al. 2015; Zhang et al. 2020a; Huang et al. 2020; Alors-Perez et al. 2021). The potential for developing intron-specific inhibitors is supported by the observation that different mammalian splicing events show different levels of sensitivity to this class of inhibitors (Vigevani et al. 2017; Corrionero et al. 2011; Finci et al. 2018; Cretu et al. 2018; Seiler et al. 2018). The sequence neighborhood surrounding and including the BP-A and polypyrimidine tract influence inhibitor sensitivity *in vitro*, in the few cases tested (Vigevani et al. 2017; Finci et al. 2018). *In vivo* estimates of splicing changes using steady state RNA level measurements are generally partly confounded by gene-specific transcription and decay rates, however short, GC-rich introns seem most affected (Vigevani et al. 2017; Seiler et al. 2018). Much about the specific intron features that might predict sensitivity remains to be learned.

We set out to study intron-specific features of splicing inhibition by splicing inhibitors using *S. cerevisiae*. Previous studies have shown that yeast and other organisms are naturally more resistant to splicing inhibitors than mammals, and have patterns of amino acid substitutions in the BP-A binding pocket of their SF3B1 homolog that account for this resistance (Serrat et al. 2019; Hansen et al. 2019; Carrocci et al. 2018). Moreover, replacing the existing amino acids with those from human SF3B1 increases sensitivity to splicing inhibitors, at least for *S. cerevisiae* and *C. elegans* (Serrat et al. 2019; Hansen et al. 2019; Carrocci et al. 2018). We used CRISPR/Cas9 to introduce 14 specific amino acid changes in the BP-A binding pocket of the yeast SF3B1 homolog HSH155, making it identical to human SF3B1 through the region of the protein that binds the BP-A and splicing inhibitors. Addition of Plad-B to this yeast strain results in rapid splicing inhibition. RNA-seq analysis of the transcriptome after treatment with Plad-B or Thailanstatin-A (Thail-A) reveals patterns of response to inhibitors that we suggest are due to intron-specific differences in the rates of mechanistic steps that lead to branch point selection. Using simple single-intron transcripts such as are common in yeast may enable more direct and parallel determination of structure-activity relationships for splicing inhibitors, possibly leading to inhibitors with improved therapeutic efficacy.

## RESULTS

### Design and creation of a humanized HSH155 inhibitor-sensitive protein

To create an allele of yeast *HSH155* sensitive to splicing inhibitors, we aligned HEAT repeats (HR) 15 and 16 from human SF3B1 and Hsh155, and identified residues that differ (Fig 1A, see also Hansen et al. 2019). Previous humanized alleles (Hansen et al. 2019; Carrocci et al. 2018) replaced HRs 5-16 from human SF3B1 in their entirety, or just 1-2 amino acids (N747 and L777) in the region of the pocket. Our goal was to replace the surface surrounding the yeast BP-A binding pocket with the human side chains, in the hope of producing a replica of the human drug binding surfaces for multiple different inhibitors, without substituting other surfaces of Hsh155 in a way that might compromise its interaction with its endogenous cofactors like Cus1, Cus2, Prp5, or Rse1. Using CRISPR/Cas9, we cleaved *HSH155* in a strain deleted for three “drug exporter” genes (*PDR5*, *SNQ2*, and *YOR1*, Jeong et al. 2007; Hansen et al. 2019), in the presence of a homologous repair fragment designed to direct the substitution of the 22 amino acids that differ between human and yeast to “humanize” HRs 15 and 16. In analyzing multiple independent colonies of edited yeast it became clear that strains with complete humanization of HRs 15-16 were invariably also heat sensitive (not shown). One strain, confirmed by DNA sequencing to be derived from partial incorporation of the rescue sequence, replaced the 14 most C-terminal yeast-specific residues with their human counterparts, and did not have a temperature sensitive growth phenotype (Fig 1A, B). Since replacement of yeast with human HRs 5-16 is not temperature sensitive (Carrocci et al. 2018), we suspect that interactions between HR15 and HR14 are compromised when the N-terminal segment of HR15 is humanized. In more carefully evaluating the utility of this serendipitous allele of *HSH155*, we found that amino acids that had not been humanized did not contribute to the binding surfaces for several different inhibitors of this class based on X-ray and cryo-EM analysis of human splicing complexes (Cretu et al. 2018, 2021, Fig S1). This allele, named *hsh155-ds* (ds, for drug-sensitive, see below), is expressed equivalently to wild type when fused to a C-terminal GFP-tag and evaluated by an anti-GFP western blot (Fig 1C). The surface of the binding pocket formed by PHF5A would be contributed in yeast by its homolog Rds3.

**Figure 1:**
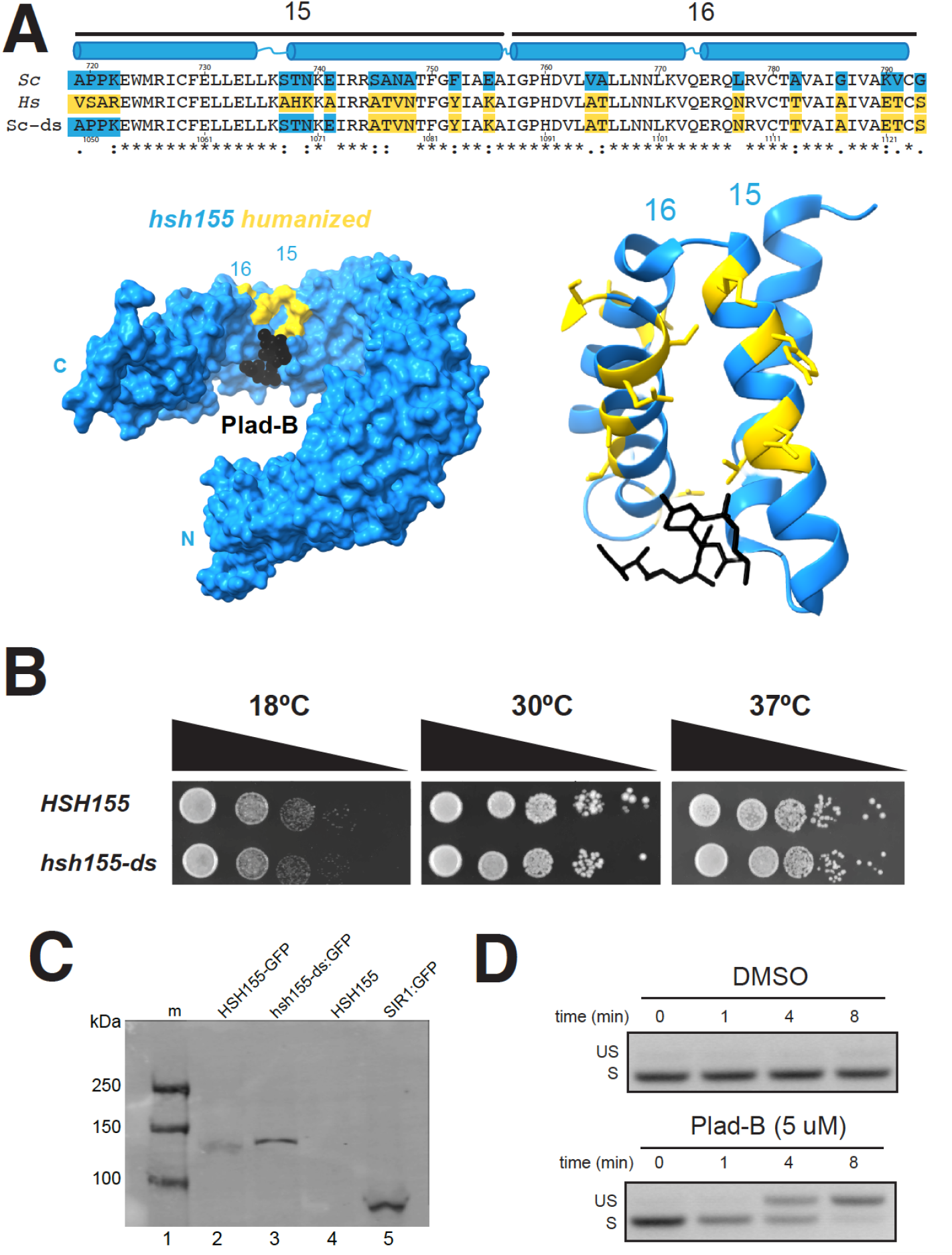
Creation and characterization of a yeast HSH155 allele humanized for sensitivity to splicing inhibitors. **(A)** Sequence alignment of the BP-A binding pockets of HSH155 (top line) and SF3B1 (middle line). HEAT repeats 15 and 16 are shown with their paired alpha helices in blue above the alignment. The bottom line shows the sequence of *hsh155-ds* with blue boxes indicating the yeast-specific amino acids, and gold boxes indicating the human-specific amino acids. Amino acids identical in yeast and humans are unshaded. The amino acid numbers are shown above (yeast) and below (human). Below the sequence is a model of SF3B1 (left) or just the HEAT repeats 15 and 16 (right) bound to Plad-B (black, adapted from Cretu et al., 2018, PDB: 6EN4). Human-specific amino acids replaced in the yeast protein are shown in gold. (**B)** Serial dilution of WT *HSH155* and *hsh155-ds* yeast strains on YPD plates to test for temperature sensitivity. **(C)** Western blot of strains expressing wild type and hsh155-ds tagged with GFP. Lane 1, markers; lane 2, WT Hsh155 C-terminally tagged with GFP; lane 3, hsh155-ds tagged with GFP; lane 4, untagged wild type Hsh155; lane 5, positive control SIR1 tagged with GFP. The blot was probed with mouse anti-GFP antibodies, visualized using secondary goat anti-mouse antibody, then scanned on a Li-Cor infrared scanner. **(D)** Measurement of Plad-B sensitivity by RT-PCR of the *MAT**a**1* first intron. Samples were taken at times after addition of DMSO (top) or Plad-B to 5 μM (bottom) at 0, 1, 4, and 8 minutes. Positions of the PCR product derived from unspliced RNA (US) and spliced RNA (S) are indicated by the labels.

There are no amino acid differences between human and yeast on the PHF5A/Rds3 surface of the pocket, thus editing of Rds3 was not necessary. We conclude that humanizing yeast in this manner does not create significant differences in growth or Hsh155 protein expression.

### Plad-B inhibits splicing in *hsh155-ds* yeast within minutes *in vivo*

To test for Plad-B inhibition of splicing in the humanized yeast strain, we treated cells with Plad-B (or DMSO carrier), isolated RNA, and analyzed splicing of the *MAT**a**1* first intron by RT-PCR. *MAT**a**1* mRNA (Miller 1984) has high synthesis and turnover rates allowing for the rapid and sensitive detection of splicing inhibition. Hansen and colleagues studied similarly modified yeast and found that growth is greatly inhibited at 5-10 μM Plad-B but cells remain viable (Hansen et al. 2019). Inhibition of splicing of *MAT**a**1* first intron is evident after 4 minutes of treatment with 5 μM Plad-B in YEPD (rich media) at 30°C, and is nearly complete by 8 minutes (Fig 1D). This experiment shows that splicing is strongly inhibited almost immediately upon addition of Plad-B to the culture, suggesting that this strain will be useful for experiments that require a strong and immediate block to splicing. This amount of Plad-B does not inhibit wild type cell growth (Hansen et al. 2019) and has no effect on splicing in wild type yeast even after an hour of exposure (see below).

### Splicing in humanized yeast extracts is sensitive to Plad-B at nM concentrations

The BP-A pocket binding class of splicing inhibitors (Plad-B, Spliceostatin-A, Herboxidiene) block splicing at or before ATP-dependent pre-spliceosome formation (Roybal and Jurica 2010; Cretu et al. 2018; Corrionero et al. 2011). To estimate the concentration of Plad-B needed to inhibit *in vitro* splicing to 50% of untreated extracts (IC_50_), we prepared cell-free splicing extracts from the humanized and wild type control strains and tested them using a radiolabeled *ACT1* pre-mRNA substrate (Fig 2A and 2B). We observe splicing inhibition (as judged by the decreased production of splicing products: first step lariat-3’ exon, second step lariat and final ligated product) at Plad-B concentrations above ∼10 nM (Fig 2A and 2C) relative to untreated *hsh155-ds* and wild type strains. We also observe inhibition of prespliceosome (A-complex) formation at similar concentrations of Plad-B *in vitro* using native gels (Fig 2B and 2D). Quantification of these gels indicates that the IC_50_ for splicing and for ATP-dependent splicing complex formation are both in the range ∼26-28 nM (Fig 2C and 2D), similar in magnitude to an adenovirus late leader-derived substrate in HeLa splicing extracts (∼100 nM, Effenberger et al. 2014). This suggests that the *hsh155-ds* strain provides a robust platform to study details of splicing inhibition by this class of compounds (see also Carrocci et al. 2018; Hansen et al. 2019).

**Figure 2:**
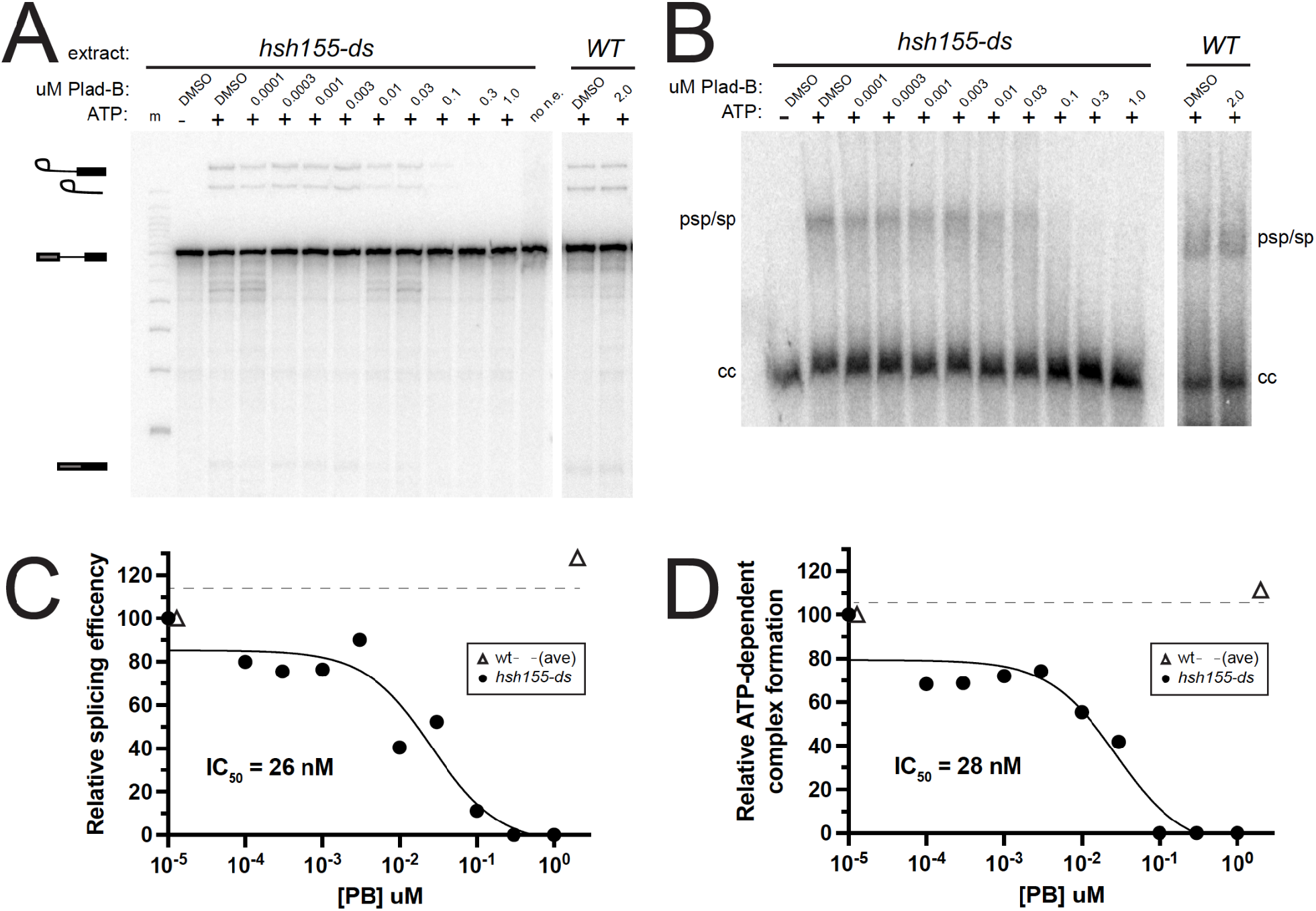
Plad-B blocks splicing during ATP-dependent pre-spliceosome formation in extracts from humanized yeast. **(A)** *In vitro* splicing assays of a 32P-radiolabeled actin pre-mRNA. Indicated extracts were incubated at 23°C for 20 minutes with either H2O or 2 mM ATP, and in the presence of DMSO or the indicated concentration of Plad-B. RNA was extracted from the reactions, resolved on a 6% acrylamide/8M urea gel and visualized by autoradiography. Lariat-3’ exon, lariat, pre-mRNA, and mature mRNA are shown. Lane “m” is a 50 bp marker and “no n.e.” is a no extract control lane. **(B)** Half of the *in vitro* splicing reactions in panel A were also directly loaded on non-denaturing gels to visualize splicing complex formation. Pre-spliceosomes/spliceosomes (psp/sp) and commitment complexes (cc) are indicated. **(C)** Quantification of splicing reactions in panel A measuring relative splicing efficiency in the presence of increasing concentration of Plad-B. **(D)** Quantification of splicing reaction assay in panel B measuring relative ATP-dependent complex formation in the presence of an increasing concentration of Plad-B.

We also tested wild type and *hsh155-ds* splicing extracts for sensitivity to the structurally distinct but mechanistically similar BP-A pocket binding inhibitor herboxidiene (HB, Hasegawa et al. 2011). We observe similar inhibition of splicing and splicing-complex formation in *hsh155-ds* extracts with IC_50_ ∼47nM and ∼74nM, respectively (Fig S2A-D). Unlike Plad-B, HB weakly but detectably inhibits *in vitro* splicing in wild type yeast extracts at the ∼2 μM level (Fig S2A-D). To test whether HB inhibits splicing *in vivo*, we constructed a reporter gene with a *MAT**a**1* first intron modified to have an open reading frame so that Neon fluorescent protein would be expressed when splicing is inhibited, and integrated it into the wild type and humanized *hsh155-ds* strains. Growth of wild type yeast for 1 hour in 5 μM HB in rich medium results in significant fluorescence above background and controls (Fig S2E), indicating splicing inhibition *in vivo* by HB. As expected the humanized strain is more sensitive, producing about 3x more fluorescence than wild type over the hour incubation (Fig S2E), consistent with the *in vitro* results here and elsewhere (Fig S2A-D, Hansen et al. 2019). These experiments show that the substitution of 14 amino acids in Hsh155 protein with the corresponding human SF3B1 residues creates a binding pocket for Plad-B (or HB), substantially increasing the ability of these compounds to inhibit splicing in yeast. The amount of inhibitor needed to block splicing *in vitro* is similar to amounts needed to block human splicing *in vitro*, with the block occurring at or near the step of prespliceosome or A-complex formation in both systems.

### Transcriptome-wide effect on splicing in the presence and absence of Plad-B and Thail-A

To examine the global impact of treatment with Plad-B and Thail-A on splicing and gene expression in yeast, we performed total RNA sequencing (RNA-seq) on wild type *HSH155* and *hsh155-ds* cells treated with with either DMSO carrier, 0.5 μM Plad-B, 5 μM Plad-B, or 5 μM Thail-A, a Spliceostatin A-like compound expected to make a covalent complex with PHF5A/Rds3 (Cretu et al. 2021; Liu et al. 2013). We chose to treat for 1 hour at 30°C to allow the effects of the drug on splicing to manifest for genes with a variety of expression rates. Duplicate cultures were treated, RNA was isolated and rRNA-depleted using DNA oligonucleotides complementary to rRNA and RNAse H (sequences in Table S3.1, see Methods), libraries were prepared and sequenced. We obtained about 1.7 billion reads across all 12 samples, mapped them, and performed splicing and differential gene expression analysis as described in Methods.

We estimated a weighted splicing efficiency (SE) for each intron (Xia 2020, Fig 3A, see Methods) resulting in measurements for 267 splicing events. Comparisons of each replicate pair (Fig S3A-F) were highly correlated, allowing us to average replicates from each treatment and use them to compare treatments to each other with confidence (Fig 3 B-E). Comparison of wild type cells treated for 1 hour with 5 μM Plad-B to those treated with DMSO show no evidence that splicing is inhibited (*r^2^* = 0.99, Fig 3B). Additionally, comparison of SEs in the wild type and humanized strain in the absence of inhibitors suggest that splicing of most introns is unchanged in the mutant (*r^2^* = 0.98, Fig 3C). There are a small number of introns that appear to be spliced slightly more efficiently in the humanized strain compared to wild type. A histogram of the change in splicing efficiency (mutant - wt) for each intron shows that most introns change <3%, and that the few that improve seem enriched for noncanonical branch point sequences (Fig S3G). We conclude that wild type yeast is completely insensitive to Plad-B, and that although the humanizing mutation may advantage introns with variant BPs (see Carrocci et al. 2018; Hansen et al. 2019), it has little effect on the overall landscape of splicing in yeast in the absence of inhibitors.

**Figure 3:**
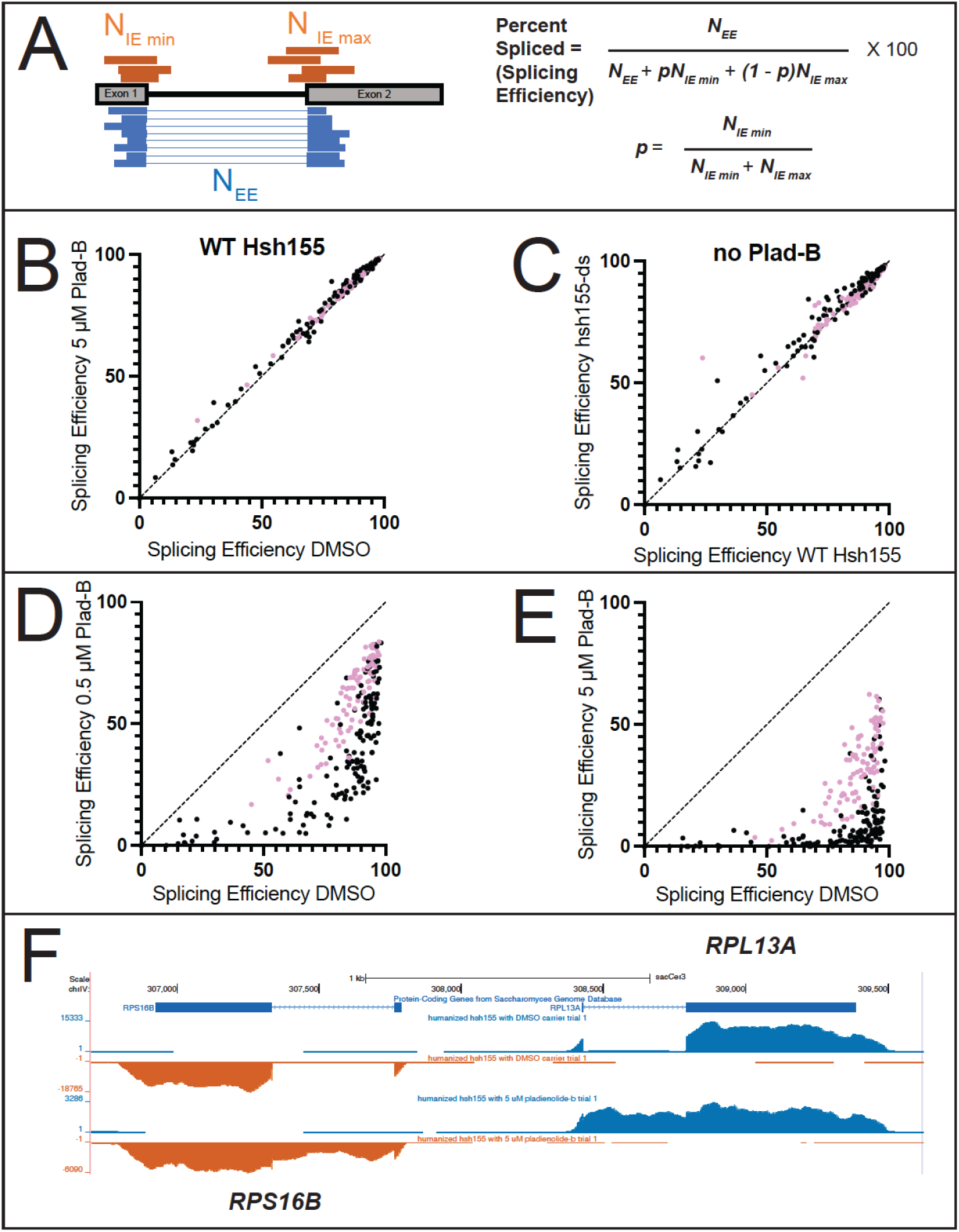
Comparisons of splicing efficiency under different genetic conditions and drug treatments. **(A)** Splicing efficiency is calculated as percent spliced using junctionCounts (see Methods). Reads spanning the splice junction (N_EE_, blue) and the unprocessed splice sites (N_IE_, orange) were used to calculate a percent spliced value as shown using the equations on the right. **(B)** Scatter plot comparing splicing efficiency for each intron in wild-type cells incubated with either DMSO (x-axis) or 5 μM Plad-B (y-axis). **(C)** Scatter plot comparing splicing efficiency for each intron in wild type cells (x-axis) or in hsh155-ds (y-axis) cells without drug (DMSO only). **(D, E)** Scatter plots comparing splicing efficiency for each intron in hsh155-ds cells treated only with DMSO (x-axis) or with 0.5 μM **(D)**, or 5 μM **(E)** Plad-B (x-axis). In panels **B-E**, pink dots represent ribosomal protein genes. **(F)** Normalized read coverage tracks over part of the yeast genome encoding divergently transcribed genes *RPS16B* (bottom strand right to left, orange) and *RPL13A* (top strand left to right, blue). Top track shows ORF locations, middle track shows humanized strain expression in DMSO (no drug), bottom track shows humanized strain expression after 1 hr in 5 μM Plad-B). Note accumulation of intron reads in the Plad-B treated sample. Browseable coverage data may be found at: https://genome.ucsc.edu/s/ohunter/Hunter_et_al_sacCer3_Yeast_Splicing_Inhibition

### Apparent splicing inhibition is Plad-B dose-dependent and different for different introns

To estimate the change in SE for each intron in the humanized yeast treated with Plad-B, we compared intron SEs from DMSO-treated cells against those treated with 0.5 μM (Fig 3D) or 5 μM (Fig 3E) Plad-B. We observe a strong, dose-dependent reduction in SE across the transcriptome of the humanized yeast strain (Fig 3D and 3E) with 5 μM showing the highest amount of retained intron for most genes (Fig 3F). None of the introns escape inhibition, but the ribosomal protein gene (RPG) introns (pink dots) appear generally less inhibited than the non-RPG introns (black dots). Many non-RPGs appear almost completely blocked after 1 hour at 5 μM (Fig 3E). Many introns spliced at 85% or more in the absence of drug show a wide range of apparent SEs (10-80%) in 0.5 μM Plad-B (see below). Because these numbers are derived from steady state measurements of spliced and unspliced RNA, it is likely that some fraction of the observed difference in intron SEs between introns are contributed by gene-specific differences in rates of transcription and decay, a concern for any analysis of splicing changes captured by measuring steady state RNA levels.

To explore the extent that gene-specific transcription and decay rates obscure potential intron-specific differences in Plad-B sensitivity, we plotted splicing efficiency as a function of Plad-B dose (Fig 4A). As suggested by the scatter plot, introns appear inhibited to different extents by Plad-B, however introns also have different apparent intrinsic splicing efficiencies in the absence of drug. To avoid complications from mRNAs with two introns or multiple alternatively spliced isoforms, we focused on single intron genes. Some but not all introns with high intrinsic (unperturbed) apparent SEs have SEs of ∼50% or more in the highest concentration of Plad-B (5 μM), while other introns that are less efficiently spliced in the absence of drug are completely inhibited at 0.5 μM (Fig 4A).

**Figure 4:**
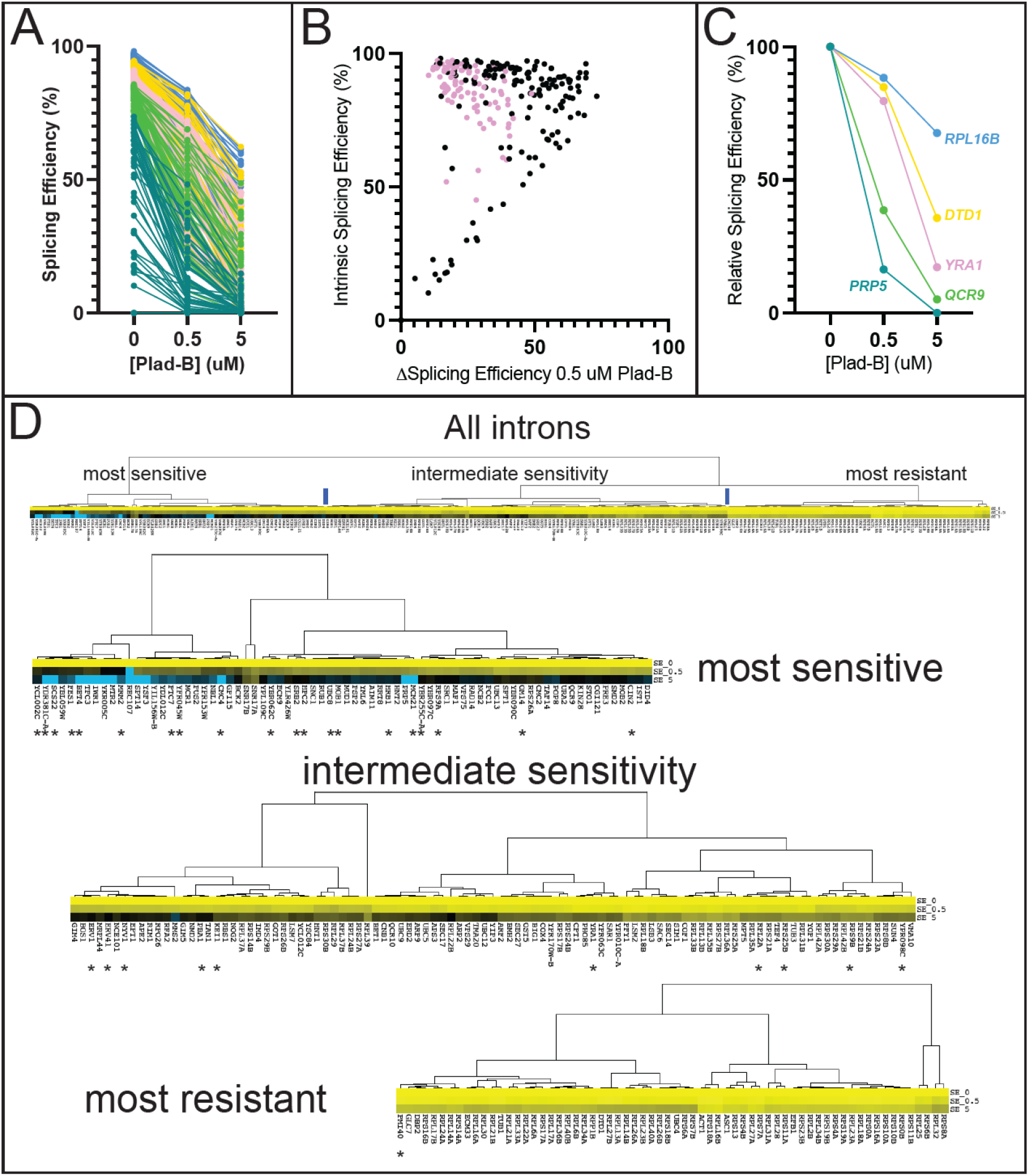
Introns can be classified by their sensitivity to Plad-B. **(A)** Splicing efficiency (SE) dose response curves for all 247 introns analyzed at 0, 0.5, and 5 μM Plad-B for the humanized strain. The data has been separated into quintiles and color-coded: Blue represents quintile 1 (most efficient), gold represents quintile 2, pink represents quintile 3, green represents quintile 4, and teal represents quintile 5. **(B)** Scatter plot comparing Plad-B sensitivity (the difference in SE between 0 and 0.5 μM Plad-B) with the intrinsic SE of the uninhibited introns (0 Plad-B, DMSO). Pink dots represent ribosomal protein gene introns. For introns >75% intrinsic splicing efficiency vs ΔSE in 0.5 μM Plad-B *r^2^* = 0.08. **(C)** Relative SE dose-response for 6 representative genes. The data shown in **(A)** has been normalized to hsh155-ds DMSO sample as described in the methods. **(D)** Hierarchical clustering of relative SE dose-response curves for single intron genes. Cluster nodes were used to call intron groups as “most sensitive,” “intermediate sensitivity,” and “most resistant.” Color indicates sensitivity to Plad-B with yellow being the most resistant, black being intermediately sensitive, and blue being the most sensitive. The top panel shows the complete set with the lower panels showing the groups. Introns with non-consensus branch point sequences are marked with asterisks.

Before bringing these introns into alignment for comparison, we asked whether there was a strict relationship between splicing efficiency in the absence of inhibitor (“intrinsic” splicing efficiency or strength) and intron-specific Plad-B resistance by plotting the change in splicing efficiency between 0 and 0.5 μM Plad-B (ΔSE0.5) against the intrinsic splicing efficiency of the uninhibited introns (Fig 4B, Supplemental Table S4.1, note: small changes in ΔSE0.5 indicate higher Plad-B resistance, points on the diagonal are introns with no apparent resistance to 0.5 μM Plad-B). As can be seen from the graph, the group of introns with intrinsic splicing efficiencies above 50% or so have a wide range of responses (10-70% inhibition) to Plad-B, whereas most of the ∼15 introns with intrinsic splicing efficiency below 50% are completely inhibited by 0.5 μM Plad-B. The intrinsic splicing efficiency of a well-spliced intron (>75%) is poorly predictive of its sensitivity to Plad-B (*r^2^* = 0.08, Table S4.2). In order to begin comparing introns with each other, we normalized their responses by adjusting their intrinsic splicing efficiency in the absence of inhibitor to 100%, and increasing the splicing efficiencies under inhibition by the same ratio, to produce relative splicing efficiency at a given inhibitor concentration for each intron (Fig 4C, for simplicity 5 introns with distinct plots are shown, Supplemental Table S4.3). Going forward, we will use the relative splicing efficiency for each intron measured after 1 hour in 0.5 μM Plad-B as a measure of response to Plad-B, the dose that best spreads out the intron-specific responses observed.

### Effects of gene expression dynamics on apparent splicing inhibition

To assess the extent to which the apparent intron-specific resistance values derived from steady state RNA measurements are influenced by gene-specific expression dynamics, we compared them to published gene-specific transcription, splicing, and expression measurements (Fig S4, Tables S4.4-4.6). Transcript synthesis rates measured by Cramer and colleagues (Miller et al. 2011) are under ∼20 molecules/cell/150min for most genes, however the RPGs produce between 50 and 400 mol/cell/150min. If splicing of all introns were completely blocked, unspliced RNA from highly transcribed genes would accumulate more rapidly and to a greater extent in 1 hour than would less frequently transcribed genes, making them appear more sensitive to Plad-B, producing a negative correlation between apparent resistance and transcript synthesis rate. However, the overall correlation is positive with an *r^2^*= 0.18, with only a few of the very most highly transcribed RPGs appearing somewhat less resistant than more slowly transcribed RPGs (Fig S4A). Thus for most genes, differences in transcript synthesis rate contribute little to apparent differences in resistance.

If all introns were completely inhibited by Plad-B, spliced mRNAs with long half-lives would disappear more slowly than would mRNAs with short half-lives, making the former appear more resistant and creating a positive correlation between apparent resistance and mRNA half life. A plot of the relative splicing efficiencies vs mRNA half life (Miller et al. 2011) shows a modest positive correlation (Fig S4B, *r^2^* = 0.42), suggesting that mRNA decay rates may partially influence estimates of resistance, making introns from more stable mRNAs appear more resistant to Plad-B. If mRNA decay rate played a large role in the apparent splicing inhibition measurement then correcting for decay would make introns appear more similar in resistance, but it does not, possibly because the intron population is dominated by RPGs that appear both more resistant as a group and have very similar half-lives. To assess the relationship between splicing rate and apparent resistance to Plad-B we used the “splicing speed” measurements of Granneman and colleagues for about 100 yeast introns (Barrass et al. 2015). Splicing speed is poorly predictive of Plad-B resistance (Fig S4C, *r^2^* = 0.06), suggesting that features governing the rate limiting step(s) of splicing for most introns in the absence of drug are different from those governing sensitivity to Plad-B. We suggest that except for the possible contribution by mRNA decay rates, the observed differences in response to Plad-B as measured by steady state RNA levels are largely due to intron-specific differences in inhibition of splicing by Plad-B.

To evaluate broad relationships in intron-specific resistance patterns, we clustered the introns by Plad-B dose-response using their relative splicing efficiencies in low (0.5 μM) and high (5 μM) Plad-B. Clustering revealed three loosely defined groups of introns separated by their sensitivity to Plad-B (Fig 4D, Tables S4.7-4.9). The node with the highest apparent relative resistance (Fig 4D, “most resistant”) is greatly enriched for RPGs, but includes other introns such as those from *ACT1* and *UBC4*. The “intermediate sensitivity” class is more heterogeneous with some RPGs at the more resistant end of the cluster, and considerable numbers of genes with diverse functions. Finally the “most sensitive” node is devoid of RPG introns but contains other essential genes. Analysis of intron features that might explain or predict sensitivity to Plad-B or other inhibitors with statistical power is limited by the small number of introns in yeast. Introns are enriched in the functional class of genes associated with ribosome function (e. g. RPGs) and intron size is bimodal, with RPG introns occupying the long (∼400 nt) class (Spingola et al. 1999). We ranked the 241 introns analyzed here from most to least resistant, as well as from longest to shortest. By their ranks, intron length and resistance are positively correlated (Spearman’s rho = 0.65, Table S4.3), however this could be confounded by unknown factors that correlate with RPG function or expression. We also evaluated the ranks of the 39 introns whose BPs deviate from UACUAAC (Table S4.3), and found that 33 of 39 have ranks below 154 of this set of 241 introns, placing the vast majority of BP variant introns in the bottom third of resistance. The highest ranking BP variant intron at rank 15 is *PMI40*, but it has two overlapping potential BPs (CACUA<u>A</u>CUA<u>A</u>C) within 11 nucleotides, possibly representing a strong BP. Ranked third is the well studied *ACT1* intron that has a consensus BP plus a strong cryptic BP upstream that plays a mysterious positive role in splicing efficiency (Kao et al. 2021). In contrast, the introns with non-standard 5’ splice sites (standard is GTATGT) are not consistently more resistant or sensitive (Table S4.3). It did not matter whether we considered the very common variant GTACGT as standard or non-standard site; the effect of 5’ss sequence on resistance appears weaker than that of the branch point. Pinpointing the physical features of introns that potentiate or depress the effect of the splicing inhibitors must await more detailed experimentation, but to a first approximation BP strength appears to be associated with Plad-B resistance.

### Intron-specific sensitivity to Plad-B manifests during co-transcriptional splicing

To measure the effect of Plad-B during co-transcriptional splicing, we used single-molecule intron tracking (SMIT), which evaluates splicing on chromatin-associated nascent transcripts (Oesterreich et al. 2016; Alpert et al. 2020). We treated the wild type and the humanized strains with 5 μM Plad-B for 15 min and processed cells for SMIT, sequencing chromatin associated nascent RNA from 63 intron-containing genes, mapping their 3’ ends to obtain the position of RNA polymerase, and determining the status of the intron on the nascent transcript (spliced or unspliced, Fig 5A, Oesterreich et al. 2016; Alpert et al. 2020). Since SMIT requires the use of gene-specific 5’ primers to capture nascent transcript reads, the number of genes analyzed was limited. However, we obtained sufficient data to graph the relationship between RNA polymerase progression and nascent transcript splicing for 37 intron-containing genes (6 are shown in Fig 5B, for complete data see Table S5). To estimate co-transcriptional splicing efficiency (CTX-SE), we calculated the area under the curve (AUC) within a defined window starting from the polymerase position at the onset of splicing to the position 200 bp downstream from the onset point, a narrow snapshot of the gene expression pathway which should allow fair gene to gene comparisons (Fig 5A). The area under the SMIT curve (fraction spliced vs pol II position) for introns in control cells was set to 1 and the AUC from Plad-B-treated humanized cells was subtracted to calculate ΔAUC, thus an estimate of cotranscriptional Plad-B resistance (CTX-ΔSE) is given by (1 – ΔAUC) x 100% (see Methods, Fig 5B, C). Because this assay measures only nascent transcripts at steady state within this narrowly defined window, it is less likely to be confounded by gene-specific transcription or decay rates. The set of 37 introns for which we could obtain cotranscriptional Plad-B sensitivity data show a wide range of responses (Fig 5C), some with less than 20% residual splicing of the nascent transcripts within the window of analysis, whereas others achieve 80% of the splicing observed in the absence of drug.

**Figure 5:**
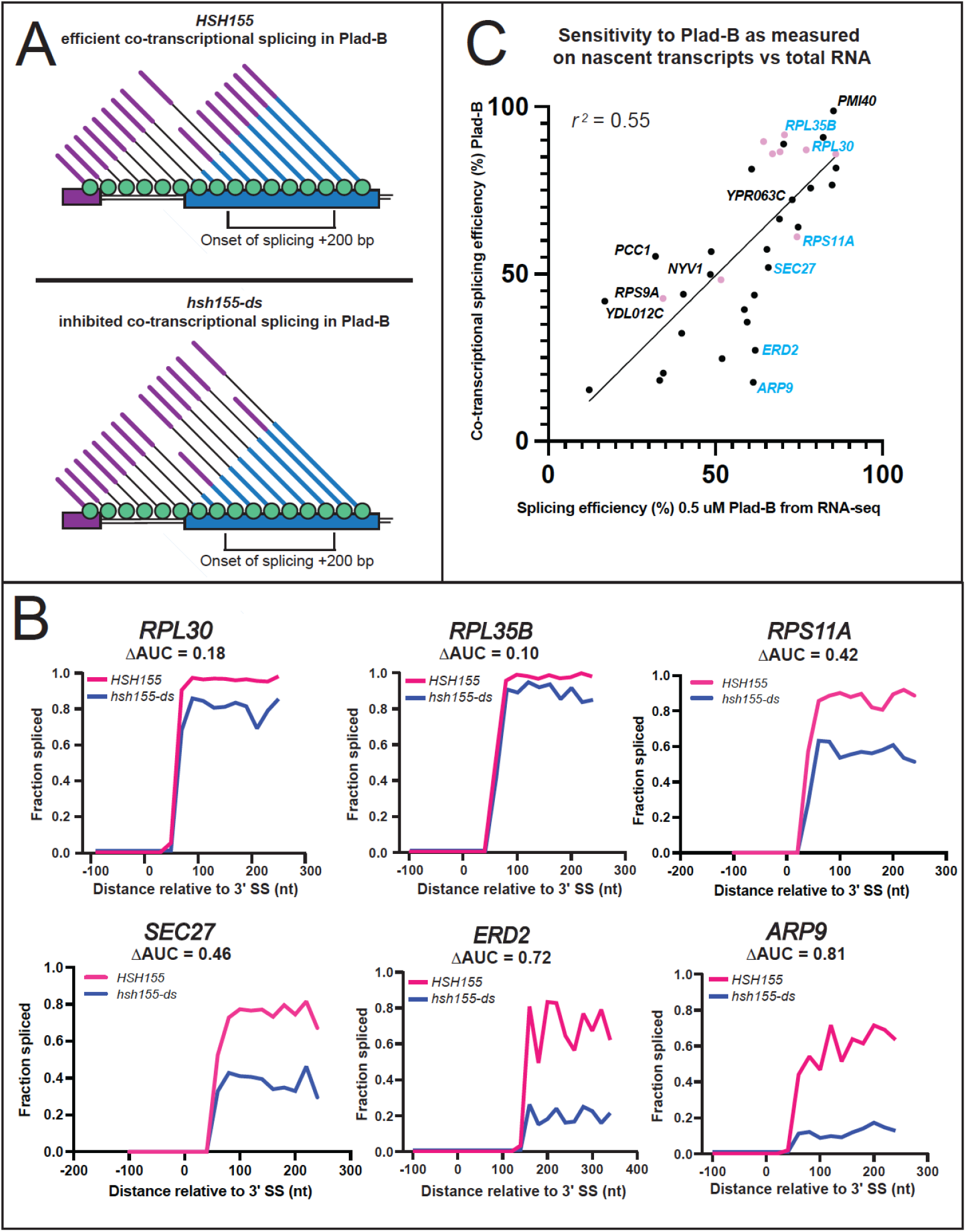
Intron-specific difference in Plad-B sensitivity during co-transcriptional splicing. **(A)** Model depicting co-transcriptional splicing in Plad-B for insensitive wild type cells (top) and sensitive humanized cells (bottom). An example gene is represented by the horizontal boxes, with RNAP II shown as green circles moving left to right during transcription. Nascent transcripts extend from RNAP II and their splicing status is shown by the presence or absence of the intron (thin line). Exon 1 is purple, exon 2 is blue. The bracket below the gene shows the range of RNAP II positions from which nascent transcripts are counted to evaluate splicing efficiency. Inhibition is detected by counting the fraction of nascent transcripts within this window that have had their introns removed. **(B)** Plots of fraction spliced versus RNAP II distance from the 3’ splice site (3’SS) for 6 genes. Pink line is WT *HSH155* with 5 μM Plad-B, blue line is *hsh155-ds* with 5 μM Plad-B. The difference of the normalized area under the curve values (ΔAUC) are shown. **(C)** Plot of splicing efficiency in 0.5 μM Plad-B measured using RNA-seq of total RNA with co-transcriptional splicing efficiency in Plad-B as measured by SMIT for 37 genes. Pink points are RPGs, genes plotted in **B** are shown in blue.

We compared the co-transcriptional estimates of Plad-B splicing resistance to RNA-seq derived estimates for the genes we could evaluate by SMIT (Fig 5C). These are reasonably similar over this subset of introns (*r^2^* = 0.55). We conclude that introns differ from each other in some fundamental way that affects (1) how well Plad-B binds the spliceosome or (2) how well bound Plad-B can interfere with their splicing, or both. Furthermore this behavior is not explained by gene-specific differences in rates of transcription, splicing, and decay. It is possible that unspliced nascent transcripts in the nucleus are subject to decay by the nuclear exosome or Rat1 nuclease at intron-specific rates (Bousquet-Antonelli et al. 2000; Egecioglu et al. 2012) after Plad-B inhibition, however such measurements have yet to be made. Some of the outliers that show higher co-transcriptional sensitivity than expected based on their RNA-seq estimated resistance, such as *ERD2* or *ARF9,* might be spliced after release from chromatin more readily than other pre-mRNAs. We conclude that a major underlying explanation for intron-specific differences in apparent sensitivity to Plad-B is that introns possess intrinsic differences that affect their inhibition by Plad-B at the spliceosome.

### Intron-specific differences in relative sensitivity to Plad-B versus Thail-A

To test whether intron-specific differences in sensitivity to Plad-B are similar to other BP-pocket binding splicing inhibitors, we compared splicing efficiency of yeast introns in 5 μM Plad-B to their splicing efficiency in 5 μM Thail-A. Thail-A is in the same class as Spliceostatin A (SSA), whose scaffold is distinct from the scaffolds of either Plad-B or HB, and which can covalently bind to a cysteine in PHF5A (Cretu et al. 2021). At 5 μM, Thail-A is overall a more potent inhibitor of splicing in the humanized yeast strains than Plad-B, however its effects are not systematically proportional across introns to that of Plad-B (Fig 6A). This is more evident in a plot of the relative splicing efficiency in 5 μM Plad-B (x-axis) against the difference in splicing efficiency between Thail-A and Plad-B (y-axis, Fig 6B). A swath of introns whose relative splicing efficiency in Plad-B is between 30 and 50% vary with respect to the difference in relative splicing efficiency between the two inhibitors over a range of from 5% (slightly more sensitive to Thail-A than Plad-B, e. g. *RPS9B*, *COF1*, and *RPS13*) to 25% (much more sensitive to Thail-A than to Plad-B, e. g. *RPL17B* and *GLC7*). Although a large number of the well-separated introns in the plot are RPGs (pink dots) that are more resistant to Plad-B than other introns, non-RPGs (black dots) also differ in their sensitivity to Thail-A relative to Plad-B. Considering that each intron should have the same gene specific transcription and decay rates in these two experiments, the differences observed are likely to be due to intron-specific, drug-specific differences in splicing inhibition. Based on intron-specific differences in inhibition by Thail-A as compared to Plad-B, we conclude that introns are differently susceptible to inhibition by chemically distinct members of the general class of splicing inhibitors that bind to the BP pocket of humanized Hsh155.

**Figure 6.**
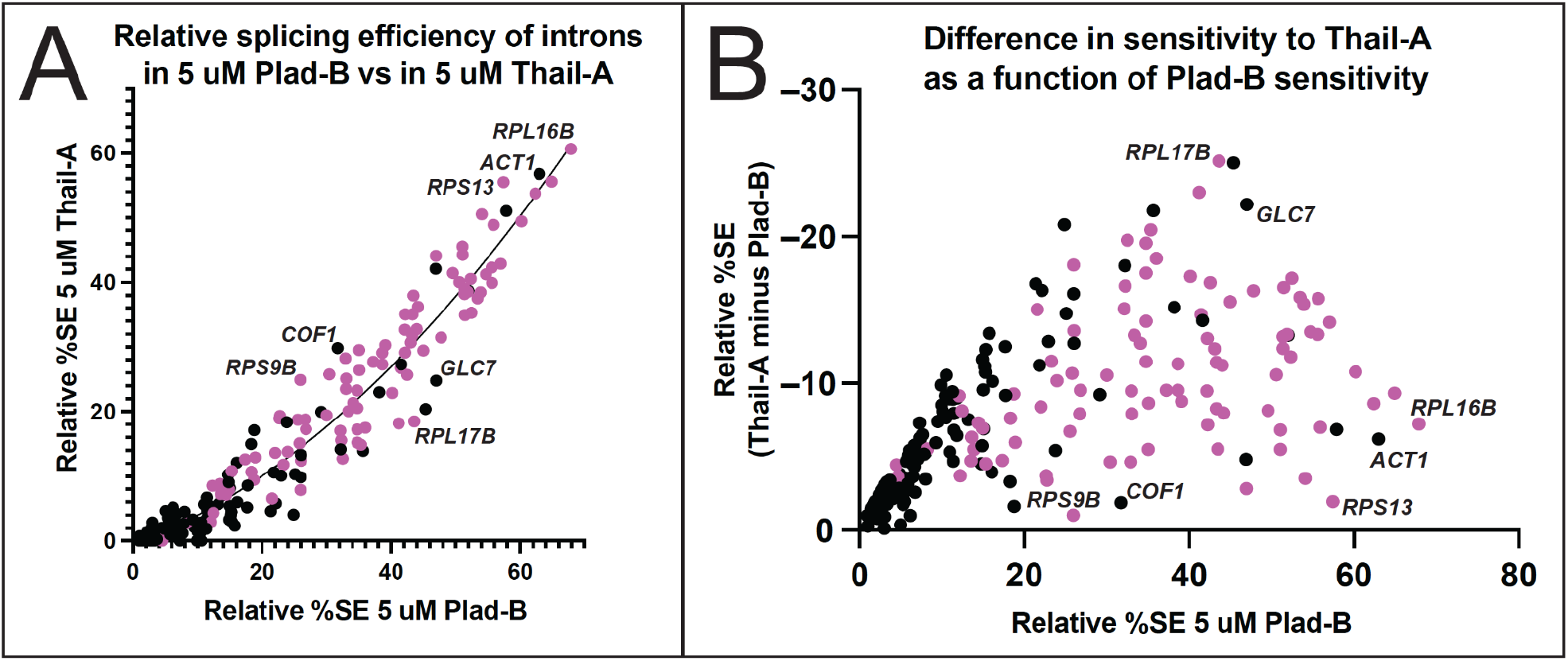
Introns respond differently to Plad-B and Thail-A. **(A)** Scatter plot comparing the relative splicing efficiency of introns in 5 μM Plad-B (x-axis) compared to their relative splicing efficiency in Thail-A (y-axis). As Thail-A is overall a more potent inhibitor than Plad-B at 5 μM all introns are more sensitive to Thail-A. The regression line is derived from a nonlinear least squares fit. Pink dots are ribosomal protein genes. Introns above this line are relatively less sensitive to Thail-A than to Plad-B as compared to the average intron, whereas those below the line are relatively more sensitive to Thial-A than Plad-B as compared to the average intron. **(B)** Relationship between sensitivity to Plad-B and the difference between Thail-A and Plad-B sensitivity. The Y-axis represents the difference between the resistance to Thail-A and Plad-B; values are negative because Thail-A is a more potent inhibitor. A high position in the Y dimension indicates an intron whose sensitivity to Thail-A is much greater than to Plad-B, whereas those lower in the Y dimension have more similar sensitivities to both inhibitors.

### Splicing inhibitors alter the expression of intronless genes

A striking feature of the *S. cerevisiae* genome is the large number of intronless genes. The existence of introns in a small subset of genes fundamental to cell function offers an opportunity to investigate the complex integration of splicing into the gene expression landscape of a eukaryotic cell by observing indirect effects of splicing inhibition on intronless genes. As observed for splicing of intron-containing genes (Fig 3B), comparison of mRNA levels for all genes in Plad-B treated vs untreated wild type *HSH155* cells showed no significant changes using DESeq2 (Love et al. 2014), indicating that Plad-B has no effect on gene expression in wild type yeast. To explore how a strong coherent block to splicing reverberates across the transcriptome, we similarly analyzed changes in mRNA levels for all genes in Plad-B treated vs untreated humanized *hsh155-ds* cells, and plotted the log_2_ fold change in expression vs the negative log_10_ of their adjusted p values using volcano plots (Fig 7A-C). As expected we observe downregulation of most intron-containing genes (Fig 7A, green points; Fig 7B), as well as numerous, presumably secondary changes in expression of intronless genes (Fig 7A, black points; Fig 7C). A few intron-containing genes appear upregulated (Fig 7B, e. g. *PCH2* and *NBL1*). Some meiotic genes like PCH2 appear transcriptionally induced (see below), and the unspliced transcripts from *NBL1* accumulate to a level about 2 fold above normal *NBL1* mRNA levels for unknown reasons, despite the block to splicing.

**Figure 7:**
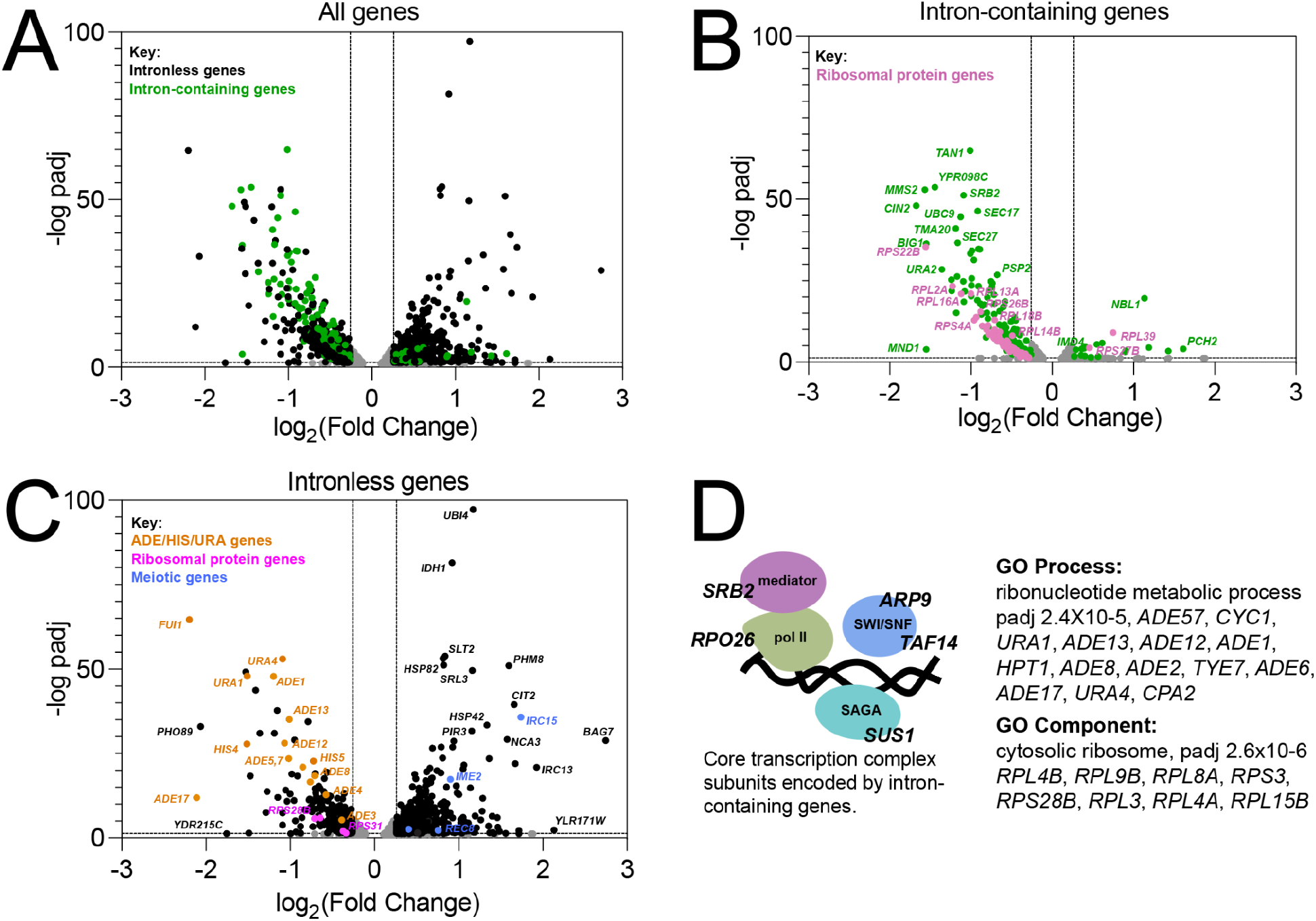
Impact of splicing inhibition on intronless genes. **(A)** Volcano plot showing differential gene expression analysis of the humanized hsh155-ds strain in 5 μM Plad-B compared to DMSO. Y-axis shows −log10 adjusted p-value and the X-axis shows log2 fold change in expression. Points not meeting p-value < 0.05 (-log padj > 1.3) or log2 Fold Change = |0.26| (Fold Change = ± 1.2) are gray. Both intron-containing (green) and intronless genes (black purple) are shown. **(B)** Same plot as in **A** but only showing intron-containing genes. RPGs are pink and non-RPGs are green. **(C)** Same plot as in **A** but only showing intronless genes. *ADE*/*HIS*/*URA* genes are in orange, RPGs are in pink, meiotic genes are in blue. **(D)** GO analysis identifies enrichment of the ribonucleotide metabolic process and the ribosome (component) in the set of downregulated intronless genes. At left is a diagram of four key protein complexes required for chromatin state and transcription along with the names of intron-containing genes encoding at least one of their subunits.

The prevalence of introns in RPGs (102 of 137 RPGs have introns) results in reduced expression of intron-containing RPGs after a 1 hour Plad-B treatment in the humanized strain (Fig 7B). This likely results in insufficient translation of ribosomal proteins to sustain ribosome biogenesis, which then feeds back on RPG transcription (de la Cruz et al. 2018; Albert et al. 2019), resulting secondarily in reduced expression of intronless RPGs (Fig 7C, pink dots). GO analysis using only intronless genes changing >1.5 fold as input confirmed that intronless RPGs are enriched in the down regulated class of intronless genes (Fig 7D). Another prominent class of down regulated intronless genes concerns nucleobase synthesis, for example numerous genes in the connected adenine (*ADE1*, *ADE5,7*, *ADE8*,…*ADE17*) and histidine (*HIS4*, *HIS5*) pathways, and some for uracil synthesis (*URA1*, *URA4*) and import (*FUI1*, orange points, Fig 7C). Although there is one intron-containing gene *IMD4* (encoding inosine monophosphate dehydrogenase, also carrying an intronic snoRNA *SNR54*) in the *de novo* purine biosynthetic pathway, expression of intronless paralogs *IMD2* and *IMD3* is unchanged, and it is unclear how loss of this enzymatic activity would repress *ADE* genes (Ljungdahl and Daignan-Fornier 2012). Mutations originally identified through their strong block to rRNA synthesis (Hartwell et al. 1970), were ultimately revealed to be in splicing factor genes (Lustig et al. 1986), likely blocking expression of intron-containing RPGs. Perhaps a sudden block to rRNA transcription produces an excess of free nucleotides that could feed back on those pathways.

At a lower threshold of 1.2 fold change, the GO analysis revealed subtle upregulation of Ty retrotransposons. Together with the apparent slight upregulation of meiotic genes (blue points, Fig 7C), the loss of expression of *ADE* genes and the upregulation of Ty elements and meiotic genes is suggestive of a general transcriptional and chromatin state defect. Considering the presence of introns in genes for at least one subunit of four main protein complexes essential for transcription (RNAP II core, Mediator, SAGA, and SWI/SNF, see Fig 7D) we suggest that general transcription is impacted within one hour of robust splicing inhibition. Depletion of chromatin remodeling and RNA polymerase II activities could lead to reduced expression of genes already induced for growth in YEPD (a poor source of adenine and uracil), and chromatin changes could reflect loss of repression of meiotic genes and Ty elements. Reduced growth efficiency might also explain the upregulation of other genes in Fig 7C that are part of stress responses and glucose exhaustion in the Plad-B treated culture.

Numerous splicing factor genes have introns, including *MUD1*, *LSM2*, *LSM7*, *SMD2*, *YSF3*, and *PRP5*, however we considered the splicing block to be so strong at the inhibitor concentrations used here that the secondary effect of loss (or gain in the case of *PRP5*) of expression of these genes would make little additional contribution to the loss of splicing activity. We did not detect a transcriptional response for splicing factor genes that would indicate the cell could sense and upregulate its splicing capacity in response to the inhibitors. We conclude that the indirect effects of splicing inhibition on the expression of the intronless class of yeast genes is complex and widespread, likely due to the importance of splicing to correct expression of the translation and transcription machinery.

## DISCUSSION

In this study we created a yeast strain sensitized to splicing inhibitors to explore the mechanism and consequences of splicing inhibition in a well-characterized splicing system. Previous work captured the effects of 4 different combinations of amino acid substitutions in plasmid-borne *HSH155* (yeast SF3B1) on Plad-B and HB sensitivity (Hansen et al. 2019; Carrocci et al. 2018). Here we constructed a chromosomal allele of *HSH155* (*hsh155-ds*) with just 14 humanizing amino acid substitutions in the pocket where Plad-B and the intron BP-A compete for binding (Fig 1, Teng et al. 2017; Cretu et al. 2018, 2021; Finci et al. 2018). The effect of the mutations in this allele (*hsh155-ds*) agree with the previous studies of Plad-B and HB sensitivity in humanized yeast (Carrocci et al. 2018; Hansen et al. 2019). Using a reporter that detects translation of unspliced RNA, we confirmed the sensitivity of wild type yeast to a high concentration of HB *in vivo* (Fig S2), that is not apparent at lower concentrations (Hansen et al. 2019). This weak natural sensitivity to HB is greatly augmented by humanizing mutations (Fig S2, Hansen et al. 2019). Sensitivity of wild type yeast to HB but not Plad-B is consistent with structure modeling that suggests these two inhibitors bind to overlapping but not identical surfaces in the BP-A binding pocket (Cretu et al. 2018). Plad-B rapidly inhibits splicing *in vivo* (Figs 1D), and IC_50_ values in *hsh155-ds* yeast splicing extracts are similar to those observed in human extracts (Fig 2, Fig S2, see also Carrocci et al. 2018).

By examining the effect of splicing inhibitors across the transcriptome, we find that individual introns have distinct Plad-B inhibition profiles (Figs 3 and 4). Intron-specific differences in sensitivity to Plad-B extend to co-transcriptional splicing as well (Fig 5), under conditions that should be agnostic to gene-specific differences in expression dynamics, with the exception of unknown features of nascent transcript decay. Furthermore individual introns have different relative sensitivities to the related but not identical inhibitors Thail-A and Plad-B (Fig 6). Finally, we assessed the effects of splicing inhibition on the expression of intronless genes (Fig 7) due secondarily to loss of expression of intron-containing genes encoding core transcription factors and ribosomal proteins.

### Transcriptome-wide patterns of splicing inhibition in engineered *S. cerevisiae*

Plad-B has no effect on splicing of any intron in wild type yeast, however every intron in the humanized *hsh155-ds* strain is susceptible to inhibition (Fig 3). Direct comparison of splicing and gene expression between two genetically different yeast cultures is noisier than comparing different treatments on aliquots of the same yeast culture due to the lack of synchrony in the diauxic shift in different cultures (DeRisi et al. 1997), however most introns appear unaffected by the humanizing mutation in the absence of inhibitor (Fig 3C). Mutation of certain residues in this part of *HSH155* subtly improve splicing of mutant branch point *ACT1* reporter introns (Carrocci et al. 2018), although similar effects were observed with human-yeast *HSH155* domain swaps that did not include HRs 15 and 16. The humanizing allele used here is mostly neutral in the absence of inhibitor, and there are hints that it subtly improves splicing of introns with noncanonical branch points (Fig S3G).

By capturing splicing inhibition at two Plad-B concentrations, we obtained simple dose-response curves for many introns that we could cluster and rank by relative resistance to Plad-B. For introns spliced at >50% efficiency in the absence of inhibitors, there is a wide range of sensitivity to Plad-B (Fig 4B), suggesting that each intron relies to a different extent on the rate(s) of the specific step(s) affected by Plad-B that together determine its overall rate of splicing. In addition, it is not possible to use a given Plad-B sensitivity to determine the sensitivity of an intron to Thail-A (Fig 6). Ribosomal protein gene (RPG) introns as a class appear to be more resistant to Plad-B and Thail-A than introns in other genes (Figs 4 and 6). The RPG introns are also characteristically longer (∼400 nt) and often contain secondary structure elements that influence splicing (Howe and Ares 1997; Rangan et al. 2023; Spingola et al. 1999; Charpentier and Rosbash 1996; Rogic et al. 2008; Meyer et al. 2011). Even within the more resistant RPG class, a range of sensitivities is evident (Figs 4, 6). The 39 yeast introns whose branch point sequences deviate from the consensus UACUA<u>A</u>C are among the most Plad-B sensitive (Fig 4D, Table S4.3), suggesting that yeast sequences including the branch point consensus contribute to the differences in sensitivity.

These observations echo findings in vertebrate cells where strong branch points are implicated as resistance features in mammalian introns (Vigevani et al. 2017; Finci et al. 2018; Corrionero et al. 2011), and are hypothesized to mediate escape and resistance (Cretu et al. 2021). A recent high resolution structure of a human A-like complex with a pre-mRNA fragment and a spliceostatin A molecule covalently attached to PFH5A provides evidence that drug binding prevents complete formation of the U2-BP helix (Cretu et al. 2021), leaving it unloaded, with SF3B1 in the open conformation. Although Thail-A, like Spliceostatin-A should be able to form the same irreversible covalent bonds with yeast Rds3 (homolog of PHF5A), we do not know how efficient this is in vivo and whether this feature contributes to intron-specific or drug-specific differences in resistance. Inhibitors may differ in their affinity for the binding pocket (or in their off-rate from specific splicing complexes) to produce overall stronger or weaker inhibition of the spliceosome, however specific features of the intron must interact with the characteristics of a particular drug-spliceosome complex to produce idiosyncratic splicing outcomes for each intron with each compound. In an investigation of Plad-B and its close derivatives, clear inhibition of specific splicing events was observed for compounds that scored weakly if at all in cell growth assays, or in an *in vitro* splicing test using a robust substrate, consistent with intron-specific activity (Effenberger et al. 2014). Combined with the surprising selectivity of a small molecule modulator of *SMN2* exon 7 5’ splice site use (Naryshkin et al. 2014), we suggest that BP-pocket inhibitors with high selectivity for specific introns and low general toxicity may exist. How intron features contribute to the activity of splicing inhibitors on individual introns will require further experimentation, but the possibility that a spliceosome operating on a given splicing substrate will be selectively druggable does not seem that remote.

### How does the block to U2-BP helix loading result in intron-specific sensitivity to inhibitors?

Yeast introns with noncanonical branch point sequences appear more sensitive to Plad-B (Fig 4D). Variant U2-BP helices may naturally be slower to load into and trigger closing of Hsh155 in the absence of inhibitor, increasing the chance that they may be recognized as “incorrect” by the Prp5-dependent discard pathway (Zhang et al. 2021; Liang and Cheng 2015; Kao et al. 2021; Xu and Query 2007). In the presence of a bound inhibitor molecule, this slow loading would be exacerbated, enhancing discard, resulting in greater sensitivity by magnifying the tendency toward discard. Conversely introns with consensus BPs in a “fast loading” context would either outrace inhibitor binding to the pocket (or respond quickly to inhibitor dissociation), or actively displace inhibitor, or otherwise delay or pass the fidelity check before the discard pathway recruits them, resulting in resistance. In this model, drug sensitivity is mediated at the molecular level by the discard pathway, such that different introns with different intrinsic rates of Prp5 release (also influenced by BP sequence context) differ in sensitivity to an inhibitor. The Prp5 fidelity check has also been linked to an escape pathway involving use of alternative BP-As by “scanning” upstream (Kao et al. 2021), however molecular details concerning the components of the discard pathway that disassemble Prp5-stalled complexes are scant.

The role of the human Prp5 homolog DDX46 in enforcing BP fidelity in human cells is unclear. Rather than stringent adherence to UACUA<u>A</u>C as observed for yeast, mammalian branch points conform to the more degenerate sequence UR<u>A</u>Y, and are thus not able to form the uniform 6 base pair + bulged A version of the U2-BP helix found commonly in yeast. Differences from yeast in the process of loading the more variable mammalian U2-BP helix into SF3B1 include the protein p14 (SF3B6, not present in yeast) that directly binds the BP-A and the N-terminal region of SF3B1 (Schellenberg et al. 2006; Tholen et al. 2022). This protein is hypothesized to stabilize the BP-A duplex and promote SF3B1 closing during loading into SF3B1 (Tholen et al. 2022; Yazhini et al. 2021), and thus could affect DDX46 recognition of authentic U2-BP helices. Instead, mammalian BP use is affected by recurrent cancer mutations in SF3B1 that reduce BP fidelity to the detriment of accurate 3’ splice site selection (Darman et al. 2015; Alsafadi et al. 2016). These mutations lie outside the BP-A binding pocket and disrupt binding of the G-patch protein SUG1P, which promotes DHX15/PRP43 disassembly of early splicing complexes (Zhang et al. 2022; Maul-Newby et al. 2022; Feng et al. 2023). Failure to disassemble incorrectly recognized BPs results in the loss of fidelity. Thus mammalian introns that may be intrinsically more susceptible to disassembly by SUG1P and DHX15 in the absence of inhibitors or mutations could also be more sensitive to inhibitors that disrupt U2-BP helix loading. Relevant to this model is the finding that certain mutations in SUG1P produce a subtle Plad-B resistance in human cells, possibly by delaying discard of a Plad-B inhibited complex to provide time for escape of the block (Beusch et al. 2023). Although we currently have only partial understanding of the factors involved in recognition and discard of complexes poised to use an incorrect BP in either system, this class of inhibitors is positioned to take advantage of BP fidelity mechanisms to create the observed intron-specific blocks to splicing.

### Are splicing inhibitors of bacterial origin driving evolution of eukaryotic microbial spliceosomes and gene architecture in the wild?

The class of splicing inhibitors that includes spliceostatins and pladienolides were originally identified as natural products with anti-tumor activity (Kaida et al. 2007). Further exploration of this class of molecules has revealed new derivatives, in particular from the ubiquitous soil bacteria *Pseudomonas aeruginosa*, *Streptomyces platensis*, and *Burkholderia sp.* (Zhao et al. 2019), which can be induced to make gram amounts of these compounds per liter of culture (Adaikpoh et al. 2023; Eustáquio et al. 2016). At the same time, evolutionary studies have noted that ancestors of present day fungi including *Saccharomyces sp.* once had intron rich genomes (Irimia and Roy 2008), with more relaxed branch point and 5’ splice site sequences, as well as factors like SF3B6 (Yazhini et al. 2021) and U2AF1 (Schirman et al. 2021), and other splicing proteins (Sales-Lee et al. 2021; Black et al. 2023) now lost in present day *S. cerevisiae*. So far, no ecological conditions under which intron loss, splice site sequence constraint, and loss of spliceosome proteins might be adaptive have been identified. It seems clear how producing a splicing inhibitor would advantage a bacterium in competition with eukaryotic microbes for limiting resources, but what evolutionary pathways might enable eukaryotes to succeed where bacteria are producing splicing inhibitors? In addition to resistance mechanisms that reduce accumulation foreign molecules in cells, or alter the inhibitor binding site, we imagine that events such as losing introns, simplification of the spliceosome by loss of protein subunits or auxiliary factors, and accumulation of intron branch point mutations that lead to stronger U2-BP helices might minimize the impact of splicing inhibitors on cell reproduction over time. As we learn more about which organisms are naturally resistant to splicing inhibitors and whether they or their ancestors naturally resided with inhibitor-producing bacteria, the possible role of bacterial splicing inhibitors in shaping the genomes of eukaryotic microbes can be assessed.

## MATERIALS AND METHODS

### Yeast strains

All strains used throughout this work are based on JRY8012 (BY4741 *MAT**a** his3Δ1 leu2Δ0 met15Δ0 ura3Δ0* with *pdr5Δ::KanMX6*, *snq2Δ::KanMX6* y*or1Δ::KanMX6*, (Jeong et al. 2007), which we refer to as “wild type” for this study. The deletion of *PDR5*, *SNQ2*, and *YOR1* are essential for allowing the splicing inhibitors to accumulate in yeast (Hansen et al. 2019; Jeong et al. 2007).

OHY001 was constructed from JRY8012 using the CRISPR/Cas9 yeast system. Construction of the CRISPR plasmid was done essentially as described (Talkish et al. 2019b), using a single plasmid p416-TEF1p-Cas9-NLS-crRNA-BaeI. This plasmid was cut with BaeI and a guide sequence DNA targeting the genomic region encoding Hsh155 HRs 15-16 region (Table S1) was inserted using HiFi DNA Assembly (New England BioLabs) to make p416-TEF1p-Cas9-NLS-crRNA-HSH155.

Co-transformation of p416-TEF1p-Cas9-NLS-crRNA-HSH155 a 1021 bp rescue DNA (Table S1) built by SGI-DNA (La Jolla, CA) carrying humanizing mutations was performed. Transformant colonies were selected on SCD – ura, and then restreaked on 5-fluoroorotic acid (5-FOA) plates to select for colonies that lost the URA-CRISPR plasmid. These colonies were subjected to colony PCR to amplify the genomic DNA coding region of HRs 15 and 16. Humanizing mutations in this region were confirmed by Sanger sequencing of the PCR product. Although the rescue synthetic construct contained humanizing mutations from Hsh155 amino acids 719 to 793, rescued clones varied with respect to which mutations were incorporated. After testing several individual clones, we selected one for which sequencing data confirmed that only amino acids 746 to 793 had changed. This specific strain OHY001 (humanizing mutations 746 to 793, Fig 1, here called *hsh155-ds*) was used in all experiments unless otherwise indicated.

### GFP-tagging

The OHY001 (*hsh155-ds*) and JRY8012 (*HSH155*) strains were transformed with a 2493 bp PCR product bearing GFP and *S. kluyveri HIS3* genes with homologous arms to the C-terminus of the DNA coding region of the *HSH155* gene. The PCR product was generated from a pYM28-kHIS3MX6 plasmid (Janke et al. 2004) using S2/S3 primers (Janke et al. 2004) adapted for C-terminal *HSH155* gene-tagging. The exact primer sequence can be found in Table S1. Transformants were selected on SCD – hist. Colony PCR was performed to confirm the integration of GFP, and the PCR product was further confirmed by Sanger sequencing. This produced strains JRY8012-*HSH155:GFP*, and OHY001-*hsh155-ds:GFP*. The Sir2-GFP yeast strain shown in the western blot analysis was a kind gift from Rohinton Kamakaka.

### Amino acid sequence alignment

Amino acid sequences for human SF3B1 and yeast Hsh155 were taken from the UCSC Genome Browser and aligned using Clustal Omega to generate the alignment shown in Fig 1A.

### Temperature-dependent growth assay

JRY8012 and OHY001 cells were cultured overnight in 5 mL YPD media at 30°C shaking at 220 rpm. Cells were spun down for 5 mins and washed with 5 mL sterile water twice. Cells were then diluted down and allowed to grow up to OD600 = 0.5. The cultures were loaded in one column of a 96-well plate and diluted by 10-fold four times. 5 μL of cells from each dilution were dropped onto YPD plates and incubated at 18℃, 30℃, and 37℃ for two days. The experiment was done in triplicate.

### Western blots

OHY001, JRY8012-*HSH155:GFP*, OHY001-*hsh155-ds:GFP*, and Sir2-GFP were grown to log phase in 30°C overnight with 220 rpm shaking. The next day, cells were harvested at OD600 = 0.5.

Whole-cell extracts were prepared by trichloroacetic acid (TCA) method exactly as described in (Gallina et al. 2015). Total protein was then mixed with 2X protein gel loading buffer and heated at 95°C for 1 minute to denature proteins. Samples were loaded onto a 4-12% SDS-PAGE gel (Bio-Rad), transferred to a PVDF membrane, and blocked with 5% low fat dry milk in 1X phosphate buffered saline (PBS) (blocking buffer). The membrane was then washed with 1X PBS with 0.1% Tween-20 (washing buffer). Next, the membrane was probed with anti-GFP antibodies in blocking buffer (primary anti-GFP mouse monoclonal, Santa Cruz Biotechnology 1:200), visualized using goat anti-mouse IRDye 680 secondary antibody in blocking buffer at 1:15,000 (Li-Cor), then scanned on a Li-Cor infrared scanner.

### Time course of inhibition

OHY001 cells were cultured overnight and diluted down to OD600 = 0.5 the next day. Plad-B was added to 5 μM and the control was supplemented with an equivalent amount of carrier DMSO. Cultures were incubated for either 0, 1, 4, or 8 minutes. At each time point, 8-9 mL cells were rapidly placed in 10 mL aliquots of 100% ethanol pre-chilled in a dry ice-ethanol bath to stop metabolism instantaneously (Koš and Tollervey 2010). Cells were pelleted and total RNA was extracted from all samples by hot phenol/chloroform extraction essentially as described (Ares 2012). Reverse transcription polymerase chain reaction (RT-PCR) was performed using primers annealing to the first intron of the *MATA1* gene (Table S1). Bands were quantified using ImageJ (NIH) and data were visualized using GraphPad Prism (v9.4.0 for macOS).

### Splicing extract preparation and *in vitro* splicing assays

^32^P-radiolabeled actin pre-mRNA was transcribed *in vitro* using the MEGAscript T7 transcription kit (Invitrogen). *S. cerevisiae* splicing extracts were prepared from the *hsh155-ds* strain OHY001 using the liquid nitrogen method as described (Stevens and Abelson 2002), except frozen cells were disrupted using a Retsch MM301 ball mill for 5 x 3 minute cycles at 10 Hz according to Jon Staley (protocol at https://voices.uchicago.edu/staleylab/retsch-ball-mill-protocol-by-the-staley-lab/, or https://www.retsch.com/knowledge-base/extract-of-saccharomyces-cerevisiae-cells-using-retsch-ball-mi <u>ll/</u>). ATP was depleted from extracts for 20 minutes at 23°C, using 1U of *S. cerevisiae* hexokinase (Sigma-Aldrich) and 16 mM D-glucose in the presence of DMSO or the indicated concentration of Pladienolide-B (Santa Cruz Biotechnology) or herboxidiene (Focus Biomolecules). After depletion of ATP, ^32^P-radiolabeled actin pre-mRNA was added to a final concentration of 0.4 nM, and standard splicing reactions were carried out in the presence or absence of 2 mM ATP at 23°C for 20 minutes as described (Ares 2013). To visualize the precursors and products of the splicing reaction, reactions were quenched in 200 µL of RNA extraction buffer (0.3 M NaOAc, 0.2% SDS, 1 mM EDTA, 10 µg/mL proteinase K) and incubated at 65°C for 10 min. RNA was extracted from the reactions using 200 µl of acid phenol (VWR), ethanol precipitated, resolved by electrophoresis on 6% acrylamide/8M urea gels, and detected by phosphorimaging. Splicing complexes were visualized by mixing splicing reactions with 2X native loading dye (20 mM Tris/glycine, 25% glycerol, 0.1% bromophenol blue, and 1 mg/mL heparin), loaded directly on 2.1% agarose gels as described (Effenberger et al. 2013) and visualized by phosphorimaging. Splicing efficiency and ATP-dependent complex formation were quantified using ImageJ (NIH) and data (Table S2) was visualized using GraphPad Prism (v9.4.0 for macOS).

### Reverse transcription Polymerase Chain Reaction (RT-PCR) and Quantification

After extraction, total RNA was subjected to DNAse treatment (TURBO DNAse, Invitrogen). First strand (FS) synthesis was performed using a 5X FS master mix composed of 0.5 μL 1X FS buffer (Invitrogen), 0.5 μL random primer mix, 5 μg RNA, and up to 7 μL of water. This mixture was heated to 95°C for 1 minute, followed by 65°C for 1 minute, then at room temperature for 1-2 minutes. All 7 μL were added to 5 μL of 1X reverse transcriptase (RTase) master mix. 1X RTase master mix included 1.5 μL 5X FS buffer, 1 μL 0.1 M dithiothreitol (DTT, Invitrogen), 1 μL 10 mM dNTPs (Thermo Scientific), 0.5 μL 40 U/μL RNAse inhibitor (RNAsIN, Promega), and 0.5 μL 200 U/μL SuperScript III RTase (Invitrogen) or water for no RTase control. The final 12 μL mixture was incubated at room temperature for 5 minutes followed by 48°C for 25 minutes. The Zymo Research DNA Clean & Concentrator-5 kit was used to purify single-stranded DNA (ssDNA). PCR was performed on ssDNA, and products were run on a 2% agarose gel. Bands were quantified from unsaturated tif files using ImageJ (NIH), and data was visualized using GraphPad Prism (v9.4.0 for macOS).

### Sequencing of rRNA depleted total yeast RNA

Libraries were created from the following six treatments of two biological replicates (12 libraries total): 1. WT Hsh155 treated with 0.05% DMSO (0 μM inhibitor), 2. WT Hsh155 treated with 5 μM Plad-B, 3. *hsh155-ds* cells treated with 0.05% DMSO (0 μM inhibitor), 4. *hsh155- ds* treated with 0.5 μM Plad-B, 5. *hsh155-ds* treated with 5 μM Plad-B, and 6. *hsh155-ds* treated with 5 μM Thail-A. Inhibitor and DMSO treatments were made by growing WT Hsh155 and hsh155-ds in 50 mL YPD to OD600 = 0.5 and splitting them into 10 mL cultures for each treatment (DMSO, 0.5 μM Plad-B, 5 μM Plad-B, or 5 μM Plad-B Thail-A. Replicates were done at separate times with different starting cultures. Total RNA was then extracted from cells as for the time course. The quality of extracted RNA was assessed using High Sensitivity RNA TapeStation (Agilent). Yeast rRNA depletion was done using the Illumina Stranded Total RNA Prep, Ligation Ribo-Zero Plus kit (Cat. no. 20040525) using RNAse H treatment, with modification. The kit provides human rRNA oligos, however, we ordered and used a substitute DNA oligonucleotide set specific for yeast rRNA (See Table S1 for sequences). The fragmentation step was skipped due to the smaller average size of yeast mRNAs compared to human mRNAs. After barcoding and amplification, libraries were sequenced using the NovaSeq 6000 system (Illumina, Inc). Reads were trimmed to 120 x 120 bp using Cutadapt. Before mapping with STAR, the trimmed reads were aligned to a set of repeat elements from the *Saccharomyces cerevisiae* genome using BOWTIE2. Approximately 6% of the reads from each sample matched repeat elements, consisting mainly of residual ribosomal RNA (including mitochondrial 15S rRNA, 21S rRNA) and yeast Ty elements (Ty1, Ty2), and were excluded from further analysis. The remaining repeat-filtered reads were mapped to the sacCer3 genome assembly with STAR version 2.5.3 (Dobin et al. 2013).

### Splicing Efficiency calculations using junctionCounts

junctionCounts 0.1.0 (https://github.com/ajw2329/junctionCounts) was first used to extract “retained intron” (RI) alternative splicing events using an annotated genome (gtf) file and the “infer_pairwise_events” command. It was then run against the dup-removed STAR alignments with the “junctionCounts” command against the 344 RI events derived above, using default parameters. The splicing efficiency (% spliced) was calculated by dividing the exon-exon junction count by the sum of the exon-exon and intron-exon count using the equations given in Fig 3A. RI splicing events with less than 100 average read counts between replicates were removed from the total normalized read count, resulting in 287 RI splicing events total (247 single intron RI splicing events and 20 double intron RI splicing events; two intron genes were not further analyzed). The relative splicing efficiency was calculated by normalizing to splicing efficiency values for the appropriate DMSO control. Regression and two-tailed Pearson correlations were calculated using GraphPad Prism v9.4.0. Data was visualized using GraphPad Prism v9.4.0 for macOS.

### Differential Gene Expression Analysis

The dup-removed STAR alignments were converted to position-by-position coverage of the sacCer3 genome assembly using the bedtools (Quinlan 2014, v2.22.1) genomeCoverageBed command. Then for each gene, the total coverage of all its exonic positions was calculated. This was divided by 240, because each aligned paired-end read in the STAR mapping covers 240 positions in the genome. The result is an estimate of the counts of reads aligned to that gene. These counts were used as inputs to DESeq2 1.30.1 (Love et al., 2014). DESeq2 uses a negative binomial distribution-based model to estimate the variance-mean relationship and perform statistical tests for differential gene expression. Differential gene expression was determined based on a log2 fold change (FC) threshold of > 0.6 or < 304 cl:884 [Edit].0.6, and an adjusted p-value (Benjamini-Hochberg correction) significance threshold of p < 0.001. The DESeq2 analysis generated a results table containing the log2 FC, p-value, and adjusted p-value for each gene (Table S7.1).

### Ranking and Clustering Intron Resistance Estimates

To compare intron resistance profiles to each other on a common scale we normalized the raw (measured) splicing efficiencies in 0.5 and 5 μM Plad-B by dividing each by the splicing efficiency in the absence of drug and multiplying by 100. This sets the resistance to 100% in the absence of drug and proportionately adjusts the two drug treated measurements. We then sorted and assigned ranks to introns based on their resistance (adjusted % splicing efficiency, 1 = most resistant) for each of the two drug concentrations and then averaged the two ranks for each intron to obtain the “average rank” (Table S4.7). Resistance profiles were clustered using Cluster3 (de Hoon et al. 2004). Clusters (.cdt and .gtr files) were visualized with Java Treeview (Saldanha 2004). The input and output files for Fig 4 are given in Tables S4.7-4.9.

### Single Molecule Intron Tracking (SMIT)

Wild type JRY8012 and humanized OHY001 cells were each grown overnight in 50 ml YPD (yeast extract, peptone, dextrose) media at 30°C with 220 rpm overnight shaking. The following day, cells were diluted to OD600 = 0.1 in pre-warmed media and grown to OD600 = 0.5.The 50 ml cultures were then split into two 25 ml cultures and treated with either DMSO or 5 μM Pladienolide-B (Plad-B) for 15 minutes to induce splicing inhibition. After treatment, chromatin was prepared essentially as described (Oesterreich et al. 2016) with additional details as outlined previously (Vo et al. 2021). Gene targeting, sequencing, mapping, and data processing were performed following the methods described in (Oesterreich et al. 2016).

All scripts for analysis and processing of SMIT data can be found on github at https://github.com/donoyoyo/SMITstuff. The raw sequencing reads for this SMIT experiment are available in Sequence Read Archive with accession number PRJNA975902. Reads were pre-processed using bbmap clumpify (version 37.90) to remove duplicate and clump reads, which could arise from PCR artifacts or sequencing errors. Next, Cutadapt (version 1.11) was used to remove adapters and filter reads based on the presence of adapters, error rate, minimum length, and minimum overlap between the read and adapter. The following parameters were used with Cutadapt: −O 8 (minimum overlap of 8 bases), −n 2 (allowing up to 2 mismatches in the overlap region), −m 23 (keeping only reads with a minimum length of 23 bases), −e 0.11 (allowing up to 11% error rate), and --discard-untrimmed (discarding reads where no adapter was found). After adapter trimming and quality filtering, the processed reads were aligned to the sacCer3 genome using hisat2-2.1.0 with the following parameters: --no-mixed (disallowing reads that align to multiple reference sequences), and --max-intronlen 10000 (setting the maximum intron length to 10000 bases to filter out potential false positive alignments).

The SMIT curves were generated using the following workflow. Only reads associated with the 63 primed genes and intronless control genes were considered. Reads are categorized by whether they are spliced or unspliced, as well as by the position of their 3’ end relative to the 3’ splice site. Splicing is calculated as the ratio of spliced count to the sum of unspliced count and spliced count (“raw fraction spliced”). The distribution and probability of insert length are determined from the data, and the fraction spliced is normalized using the position, insert length, and lengths of the gene products (spliced and unspliced), as well as the insert length probability function as originally described in (Oesterreich et al. 2016) using the SMIT script provided in the github link above.

To quantify the splicing efficiency for each gene in each sample before comparing splicing with and without Plad-B, we defined a specific area under the splicing curve (AUC) over comparable gene positions where co-transcriptional splicing is robust in untreated cells for most genes. This specific area was bounded on the 5’ side by the gene-specific point of onset of splicing (the polymerase position at which spliced nascent transcripts first begin to accumulate) and on the 3’ side by the position 200 nt downstream of the point of onset of splicing. This definition measures cotranscriptional splicing over equivalent regions of each gene and is insensitive to differences in gene structure and activity such as absolute distance between the 3’ss and the point of onset, or the length of the second exon (all of which extend beyond the 200 bp window). Files showing the fraction spliced vs RNA polymerase II position for all genes in all conditions are given in the supplementary materials and in Tables S5.

To quantify the difference in splicing efficiency between conditions for a given gene, the fractional difference in AUC values (ΔAUC) within the 200 nt beyond the onset of splicing was determined by subtracting the AUC of one sample (for example *hsh155-ds* in Plad-B) from the AUC of another sample (for example *hsh155-ds* in DMSO), and then taking that value as a percentage to estimate the % splicing efficiency relative to control for a given gene under a given treatment. To assess the significance of the ΔAUC, Wilcoxon paired p-values were calculated. The results of these calculations are shown in Tables S5.

To obtain the best estimate of the average splicing efficiency under Plad-B treatment derived from multiple samples we considered six pairs of sample comparisons, three of which compare treatments for which the standard RNA-seq analysis showed no significant splicing changes (HSH155+Plad-B vs HSH155+DMSO, HSH155+Plad-B vs hsh155-ds+DMSO, and HSH155+DMSO vs hsh155-ds+DMSO), and three of which were expected based on standard RNA-seq to reveal splicing inhibition by Plad-B (hsh155-ds+Plad-B vs hsh155-ds+DMSO, hsh155-ds+Plad-B vs HSH155+DMSO, and hsh155-ds+Plad-B vs HSH155+Plad-B). The last three comparisons represent the hsh155-ds strain treated with Plad-B (in which splicing is inhibited) in the numerator compared with three different samples in which Plad-B is either absent (vs either strain in DMSO) or inert (vs HSH155 in Plad-B) in the denominator. Thus the best estimate for percent co-transcriptional splicing under Plad-B treatment would be represented by the average of these three comparisons (Table S4.3).

### Gene Ontology (GO) Analysis

To identify Gene Ontology (GO) terms associated with differentially expressed genes, we utilized the Saccharomyces Genome Database (SGD) Gene Ontology Term Finder tool (version 2023-05-10). First, we obtained the differentially expressed genes 1.2X fold change p< 0.05 (low stringency) and 1.5X fold change p < 0.01 (high stringency) from the DESEQ2 hsh155-ds DMSO versus hsh155-ds 0.5 μM Plad-B analysis. For all three SGD ontologies (process, function, component), we obtained GO terms for both up and downregulated genes using a background set of intronless genes. These are shown in Table S7.

### Data Deposition

The raw reads for the rRNA depleted yeast total RNA-sequencing experiment are available in the Sequence Read Archive (SRA) with accession number PRJNA972180. Sequencing data for the SMIT experiment are at the Sequence Read Archive, accession number PRJNA975902.

Publicly-available UCSC Genome Browser coverage tracks for the samples can be accessed here: https://genome.ucsc.edu/s/ohunter/Hunter_et_al_sacCer3_Yeast_Splicing_Inhibition.

## Supporting information

Supplemental_Information_Hunter_etal

## ACKNOWLEDGEMENTS

This work was supported by NIH grants R01 GM040478 (including a Diversity Supplement in support of OH) and R35 GM145266 to MA. JQC was supported by a Jane Coffin-Childs postdoctoral fellowship and a UC President’s Postdoctoral Fellowship. Thanks to Jessie Lopez Suzuki for initiating this study during her rotation. We are very grateful to Karla Neugebauer for generously hosting us in her lab to learn the SMIT protocol and in particular to Tara Alpert for detailed assistance with the analysis. Many thanks to Sarah Hansen, Aaron Hoskins, Suzanne Mays, and Juan Valcarcel for communicating unpublished results, and to Irene Beusch, Aaron Hoskins, Suzanne Mays, and Juan Valcarcel, and anonymous reviewers, for helpful comments on early versions.

