## Supplemental_Information_Hunter_etal for "Broad variation in response of individual introns to splicing inhibitors in a humanized yeast strain": Hunter_etal_FigS3G.pdf

# G

Effect of humanizing mutation on splicing efficiency of individual introns in the absence of splicing inhibitor

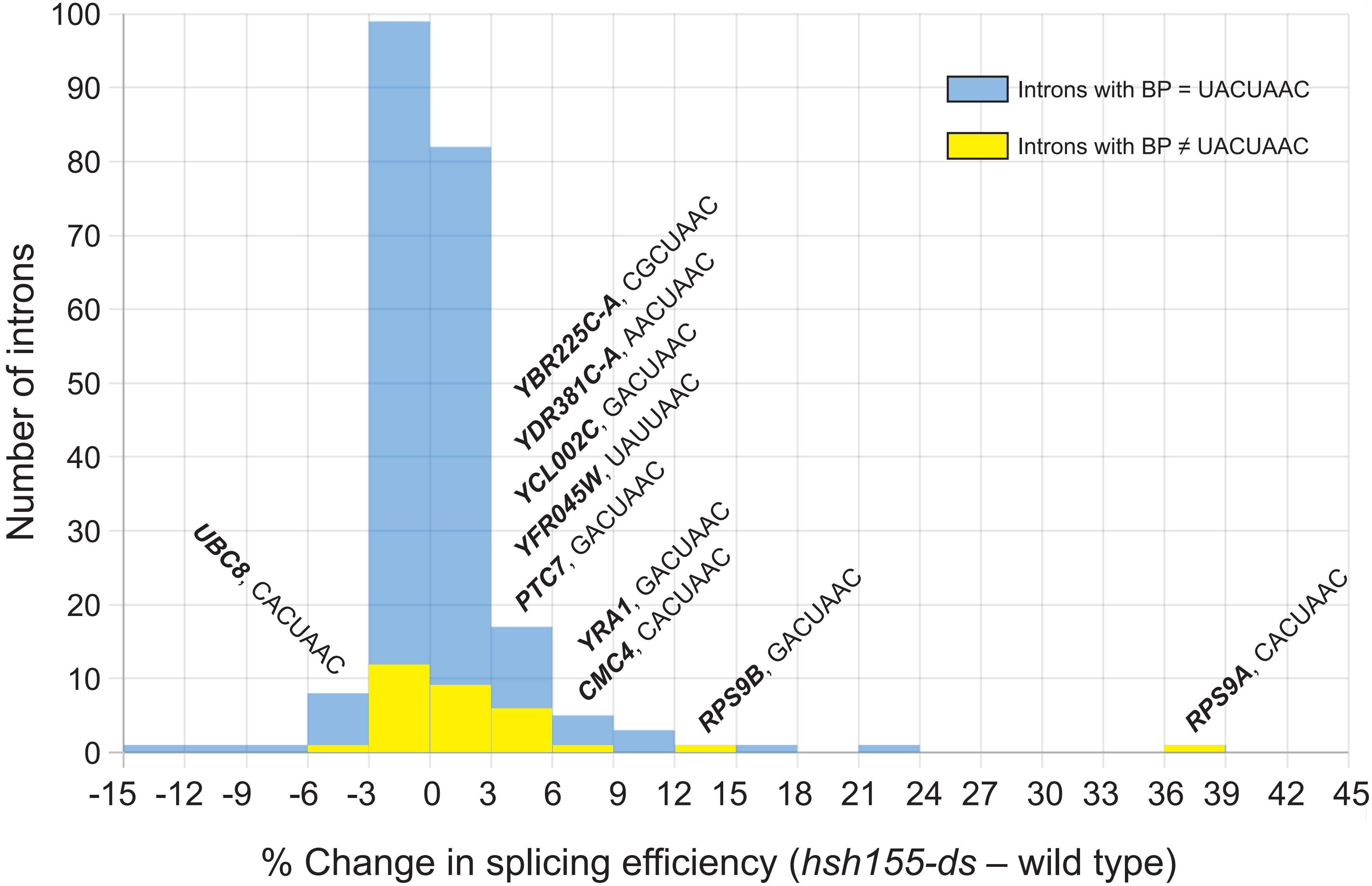


Hunter et al (Ares) Fig S4G
