## Supplemental_Information_Hunter_etal for "Broad variation in response of individual introns to splicing inhibitors in a humanized yeast strain": Supplement_Materials_Hunter_etal.docx

**Relevant to Fig 1:** Strains and sequences are found in Suppl_Tables_S1.xlsx.

**Relevant to Fig 1:** Fig S1

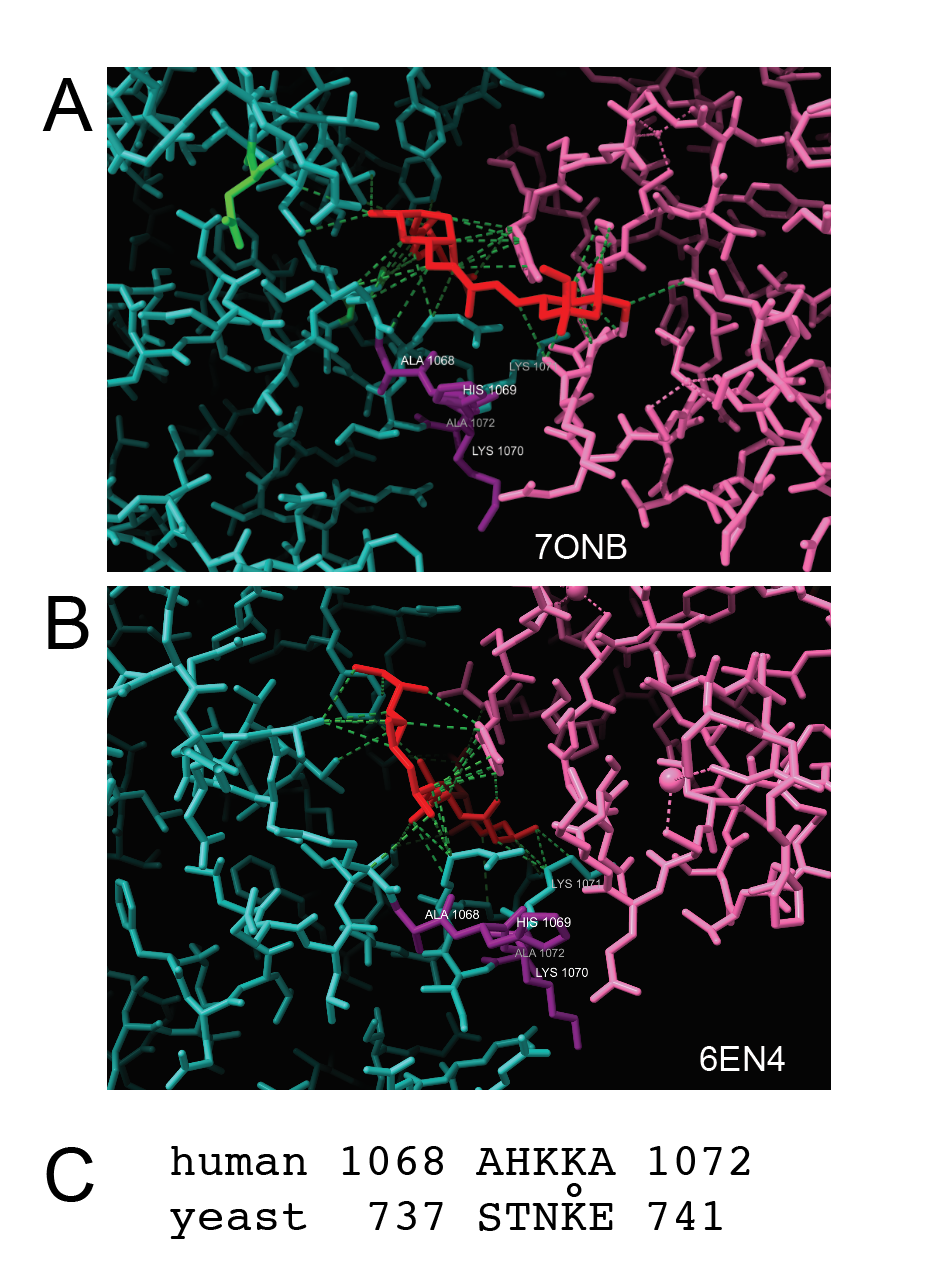

**Figure S1. Position of human residues in HR15 not replaced in *hsh155-ds*. (A)** Cryo-EM model of a human A-like complex bound with spliceostatin (7ONB, (Cretu et al. 2021). Residues originally targeted for editing but not edited are shown in purple, SF3B1 is light green, PHF5a is pink, spliceostatin is red. Protein functional groups within contact distance to the spliceostatin (~ 4Å) are connected by dashed green lines. None of the residues in purple (A1068, H1069, K1070, A1072) are within contact distance of the compound. K1071 is conserved between humans and yeast. **(B)** X-ray structure model of three SF3 protein subunits bound with a Plad-B analog (6EN4, (Cretu et al. 2018). Residues originally targeted for editing but not edited are shown in purple, SF3B1 is light green, PHF5a is pink, spliceostatin is red. Protein functional groups within contact distance to the spliceostatin (~ 4Å) are connected by dashed green lines. None of the residues in purple (A1068, H1069, K1070, A1072) are within contact distance of the compound. K1071 is conserved between humans and yeast. Note that conserved K1071 (in the same green as the rest of SF3B1) might make several contacts with Plad-B, the side chains of the other residues (in purple) point away and do not contribute to the drug binding surface. **(C)** Different sequences for this region in human (top) and yeast (bottom) at the base of the turn between the helices of HR15.

**Relevant to Fig 2:** Image analysis and calculations for quantification of Figs 2 and S2 are found in Suppl_Tables_S2.xlsx

**Relevant to Fig 2:** Fig S2

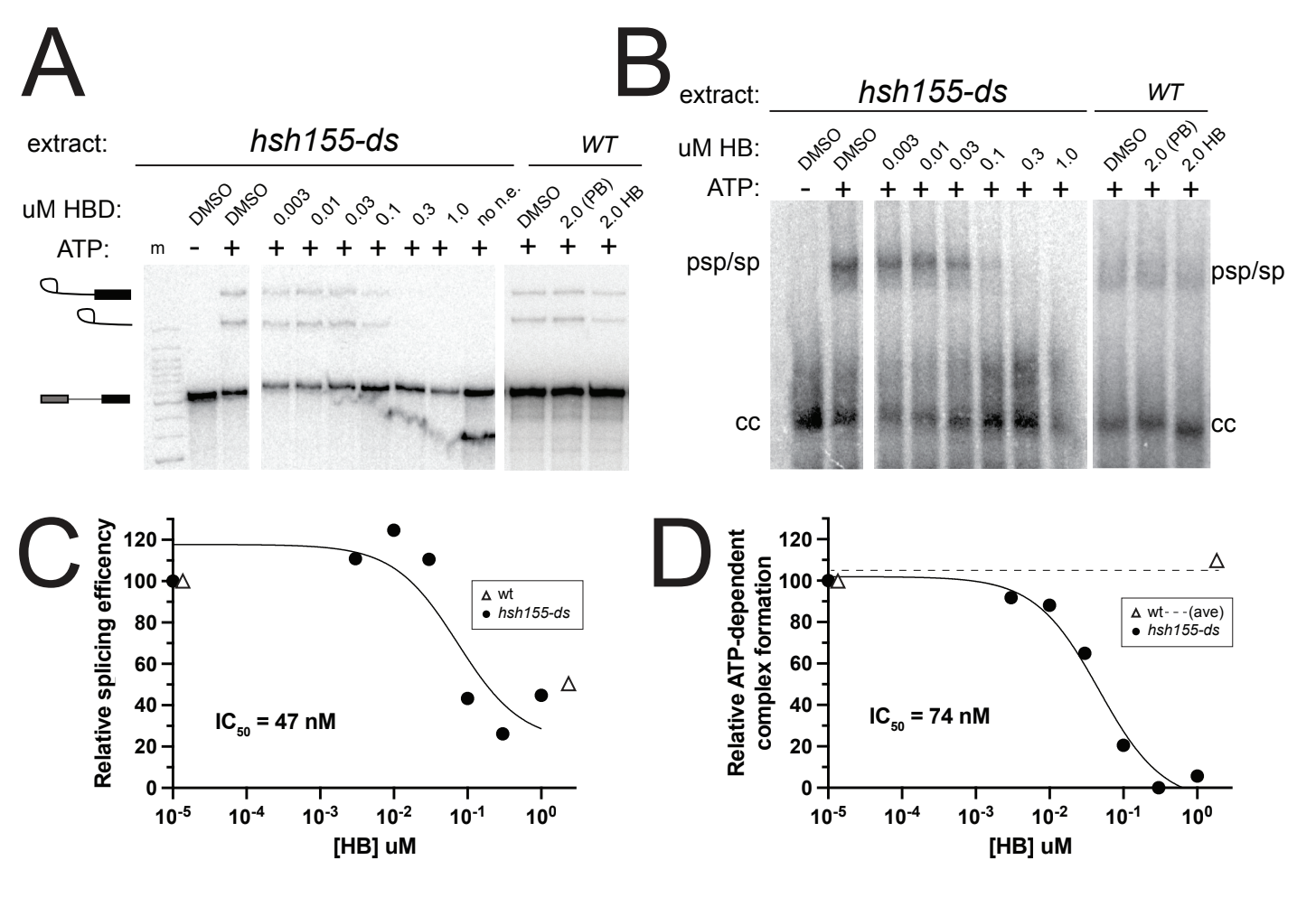
E
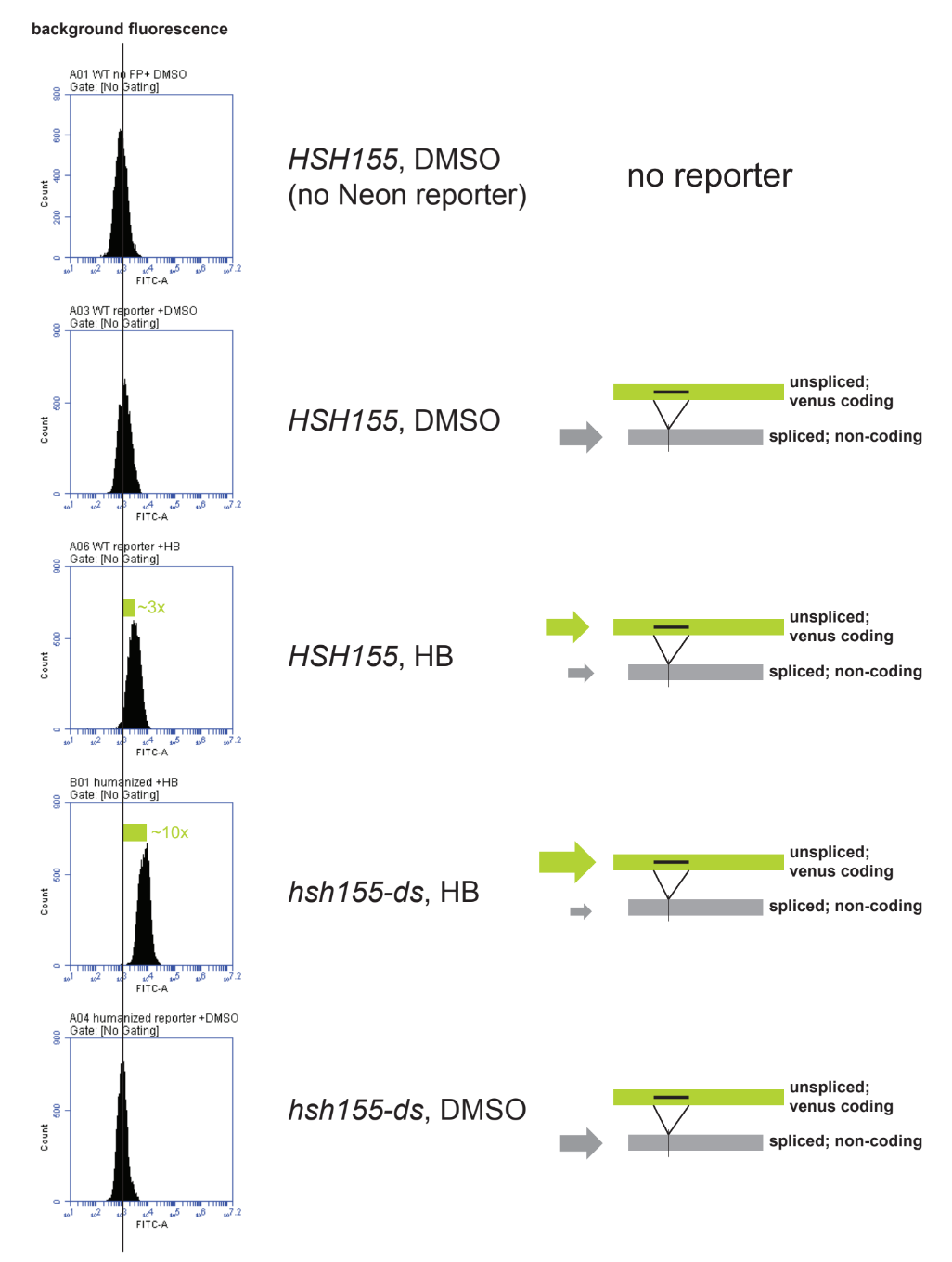

**Figure S2. Sensitivity of wild type and humanized yeast to Herboxidiene. (A)** In vitro splicing of *ACT1* pre-mRNA in extracts from humanized and wild type yeast with different concentrations of HB. Radioactive splicing intermediates and products are separated on a denaturing gel and detected with a phosphorimager. **(B)** In vitro splicing of *ACT1* pre-mRNA in extracts from humanized and wild type yeast with different concentrations of HB. An aliquot of the reaction in (A) was taken before RNA purification to preserve integrity of splicing complexes and run on a native gel. Migration of commitment complex (cc) and prespliceosomes and spliceosomes (psp/sp) are indicated on the right. **(C)**, **(D)** Graphical representation of the data extracted from the gels in **A** and **B**, respectively. **(E)** Flow cytometry of yeast strains carrying a splicing inhibition reporter that expresses neon fluorescent protein when splicing is inhibited, treated with herboxidiene.

**Supplemental Methods for the experiment in Fig S2E.**

Plasmid p1C-fl6h-sploof-venus was constructed using plasmids and methods from Dueber et al. (Lee et al. 2015) using an intron fragment designed to carry an open reading frame into the Venus coding region unless it was spliced out. DNA from this plasmid was cleaved with NotI and transformed into JRY8012 and OHY001, selecting for integration at the URA3 locus. Strains indicated in Fig S2E with or without the Venus reporter were grown to early log phase (OD 600 = 0.1) at 30°C in an Erlenmeyer flask, at which time 5 ml aliquots were transferred to glass test tubes, and incubated with 5 uM Herboxidiene, or a control amount of DMSO equivalent to that added with the compound in the treatment samples, at 30°C, with shaking. After 1 hour, 1.5 mL samples were immediately cooled in an ice-water bath (~5 min), and then centrifuged at 4°C, 2000 x g for 2 min. Supernatant was removed and pellet was resuspended in 200 uL of 2% paraformaldehyde. The cells were fixed on ice for 1 h and then centrifuged again at 4°C, 2000 x g for 2 min. The paraformaldehyde was removed and the fixed cell pellet was resuspended in 200 uL cold PBS. The ﬁxed cells were stored at 4°C. Flow cytometry was done on a BD Accuri CS6 Plus flow cytometer. Venus fluorescence was monitored in the FITC-A channel. Approximately 10,000 cells were measured and fluorescence was plotted on a logarithmic scale. The vertical line in Fig S2E represents the modal background fluorescence from yeast cells not carrying the Venus reporter. The annotated sequence of the reporter is given as a Genbank file in Suppl_Tables_S1.xlsx.

**Relevant to Fig 3:** Tables S3

**Supplemental Tables S3** are found in the file Suppl_Tables_S3.xlsx

Table S3.1 has oligo sequences recommended by Illumina for depletion of yeast rRNA for library preparation.

Table S3.2 has junction counts and splicing efficiencies for single intron genes

**Relevant to Fig 3:** Fig S3

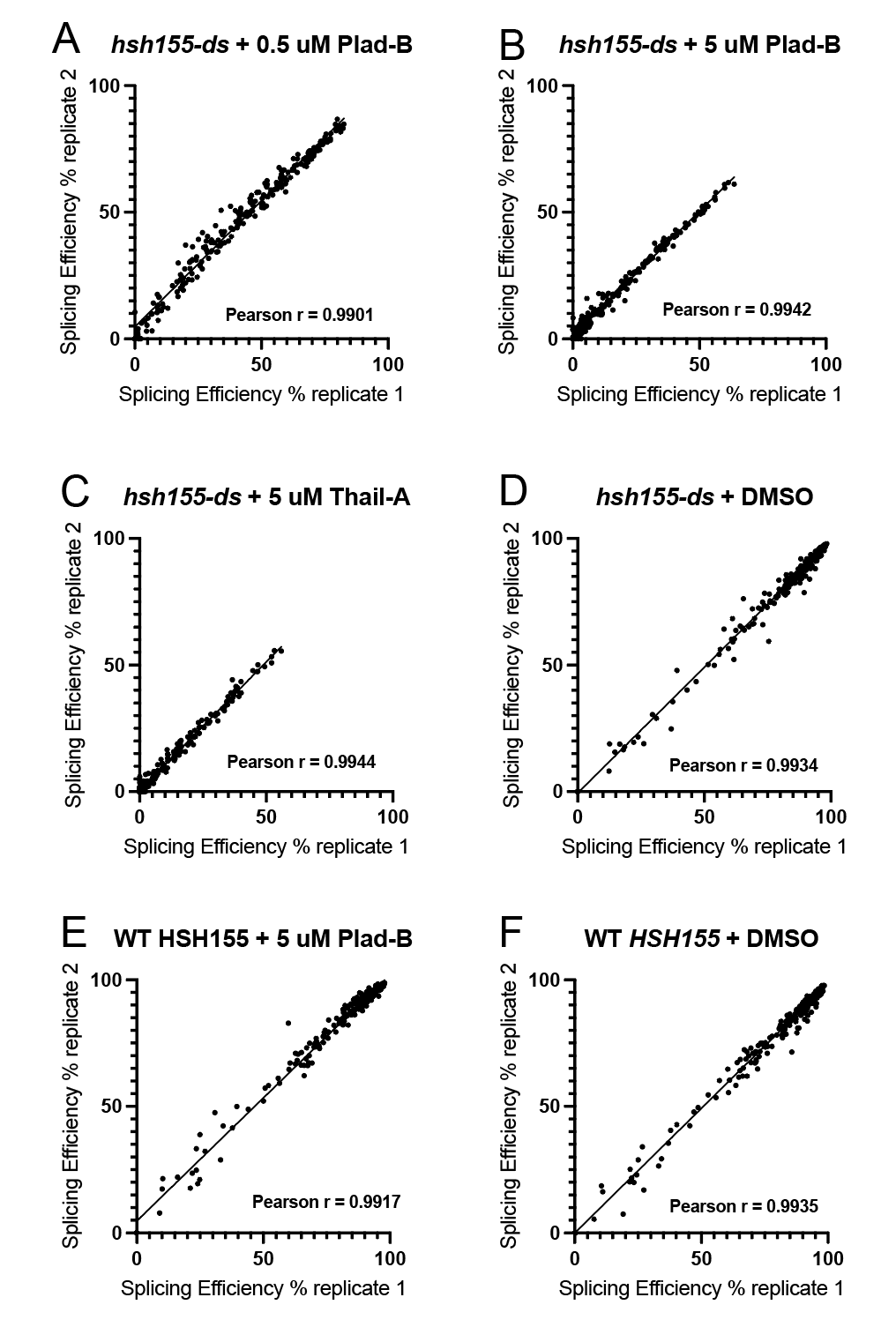

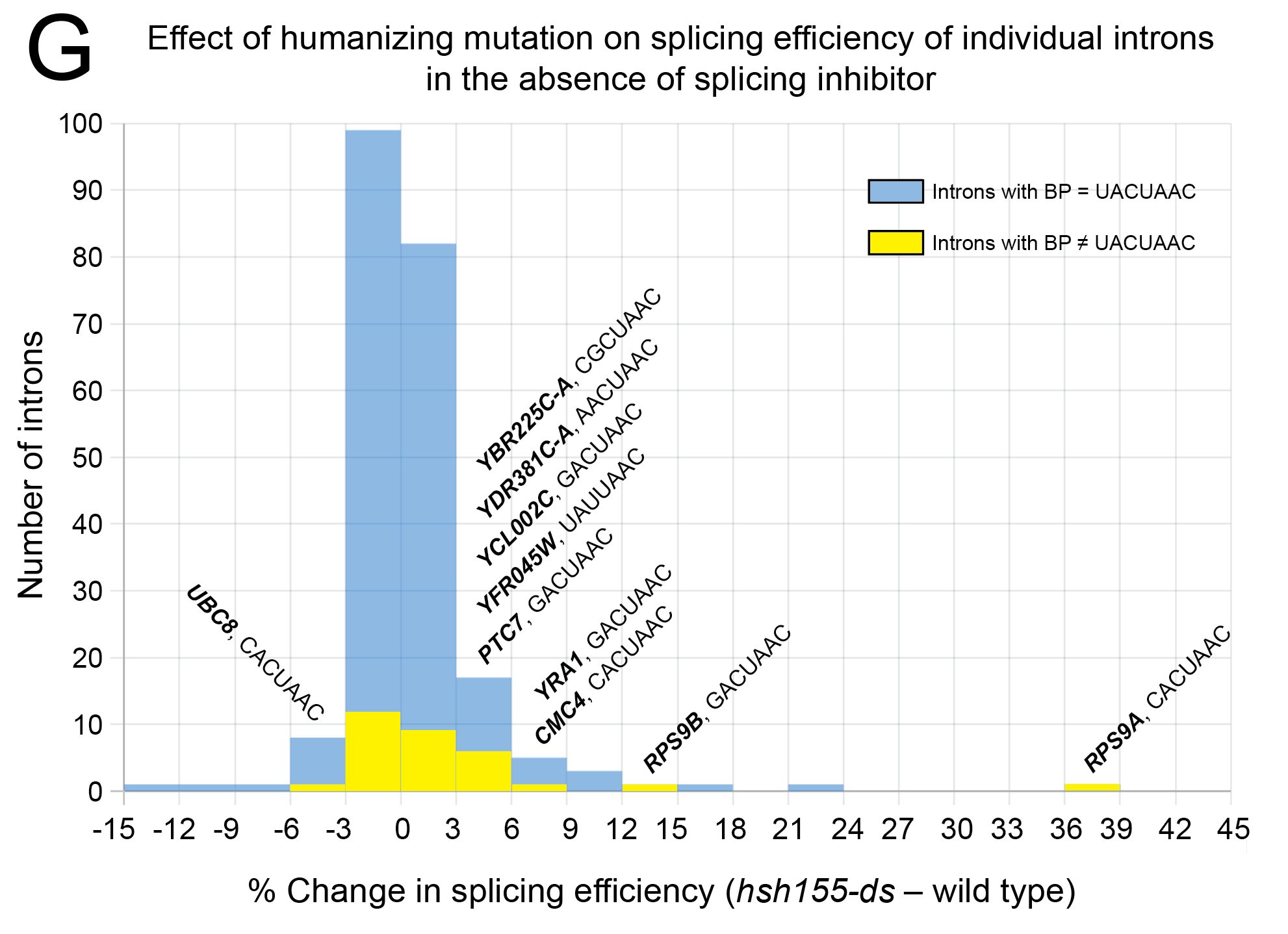

**Figure S3. Comparison of replicates and evidence the humanizing mutation subtly affects splicing in the absence of inhibitors. (A-F)** Splicing efficiency was calculated as described in the text separately for each sample and replicates were compared using scatter plots. Based on the high correlation between replicates shown here, sample replicates were pooled for comparisons between treatments. Replicate correlation plots of the six duplicate RNA-sequencing libraries. Each plot shows the Splicing Efficiency (%) of replicate 1 plotted vs replicate 2. The Pearson r for each plot is shown. **(G)** Comparison of splicing in wild type with humanized yeast in the absence of inhibitor suggests small but significant differences (Fig 3C, main text). Histogram of these differences suggests improvement of splicing for some introns in particular those with noncanonical branch points (see text).

**Relevant to Fig 4:** Tables S4

**Supplemental Tables S4** are found in the file Suppl_Tables_S4.xlsx

Table S4.1 represents the data plotted in Fig 4B.

Table S4.2 compares intron-specific intrinsic splicing efficiency with response to Plad-B.

Table S4.3 shows many aggregated intron features and measurements for each intron.

Table S4.4 represents the data plotted in Fig S4A, relative splicing efficiency (0.5 uM) versus RNA synthesis rate (Miller et al. 2011).

Table S4.5 represents the data plotted in Fig S4B, relative splicing efficiency (0.5 uM) versus RNA half-life (Miller et al. 2011).

Table S4.6 represents the data plotted in Fig S4C, relative splicing efficiency (0.5 uM) versus splicing speed (AUC) (Barrass et al. 2015).

Table S4.7 shows relative splicing efficiency data table input into clustering analysis in Fig 4D.

Table S4.8 is the output cdt file of the clustering analysis in Fig 4D.

Table S4.9 is the output gtr file of the clustering analysis in Fig 4D.

**Relevant to Fig 4:** Fig S4

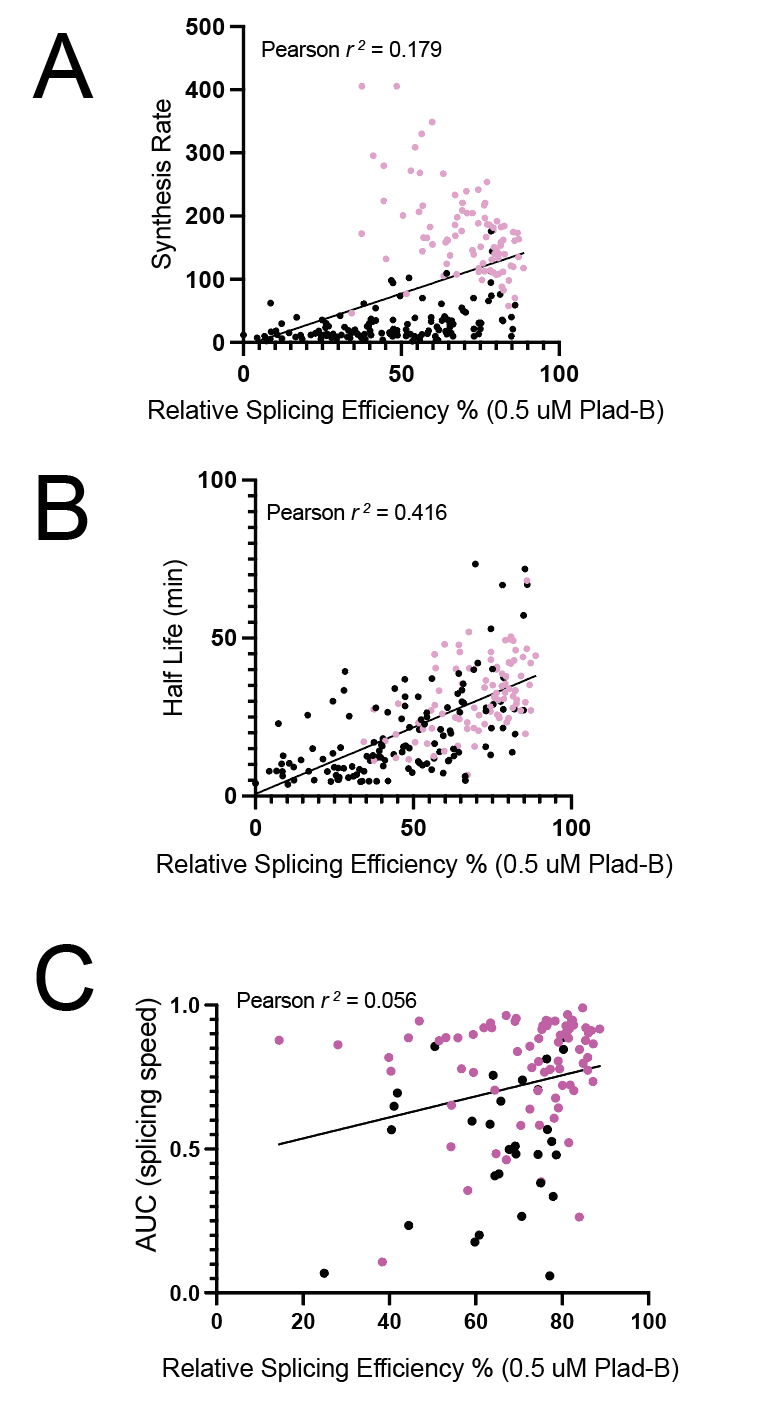

**Figure S4: Plots comparing Plad-B sensitivity (%SE0.5P) to gene-specific rates of synthesis, decay, and splicing.** **(A)** Splicing efficiency in Plad-B vs mRNA synthesis rate, **(B)** Splicing efficiency in Plad-B vs mRNA half-life. **(C)** Splicing efficiency in Plad-B vs “splicing speed”. Magenta points are ribosomal protein genes and black points are non-ribosomal protein genes. See text.

**Relevant to Fig 5:** Tables S5 and file csvs.zip

**Supplemental Tables S5** are found in the file Suppl_Table_S5.xlsx

File csvs.zip contains .cvs files from SMIT analysis representing fraction spliced vs 3’ end position for tested introns under each of 4 conditions, plus a README file.

Suppl_Table_FigS5.xlsx contains several sheets.

The first six hold the csv files used to make the plots in Fig 5B.

The next six sheets have area under the curve analysis of the csv files in all pairwise experimental comparisons, including plots of the comparisons.

The last sheet represents the data plotted in Fig 5C.

**Relevant to Fig 6:** Table S6

**Supplemental Table S6** is found in the file Suppl_Table_S6.xlsx

Table S6 represents the data plotted in Fig 6A and B.

**Relevant to Fig 7:** Tables S7

**Supplemental Tables S7** are found in the file Suppl_Tables_S7.xlsx

Table S7.1 represents the data plotted in the volcano plots in Fig 7.

The remaining sheets represent the output of the gene ontology (GO) analysis summarized in Fig 7D.
