## Supplementary figures and images for "Broad variation in response of individual introns to splicing inhibitors in a humanized yeast strain"

### Hunter_etal_FigS1.pdf

A

B

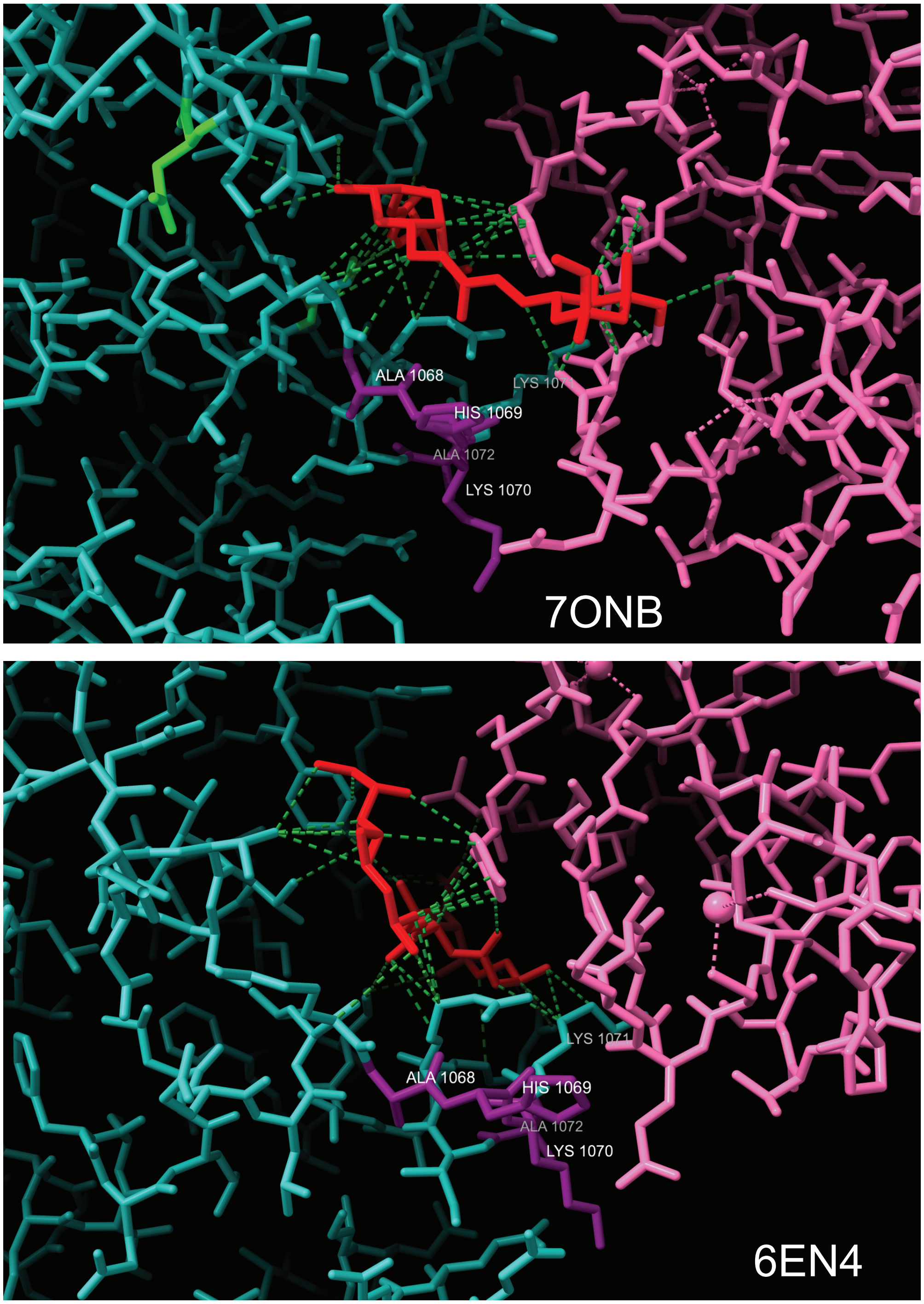


C

```
human 1068 AHKKA 1072
              °
yeast  737 STNKE 741
```

### Hunter_etal_FigS2A-D.pdf

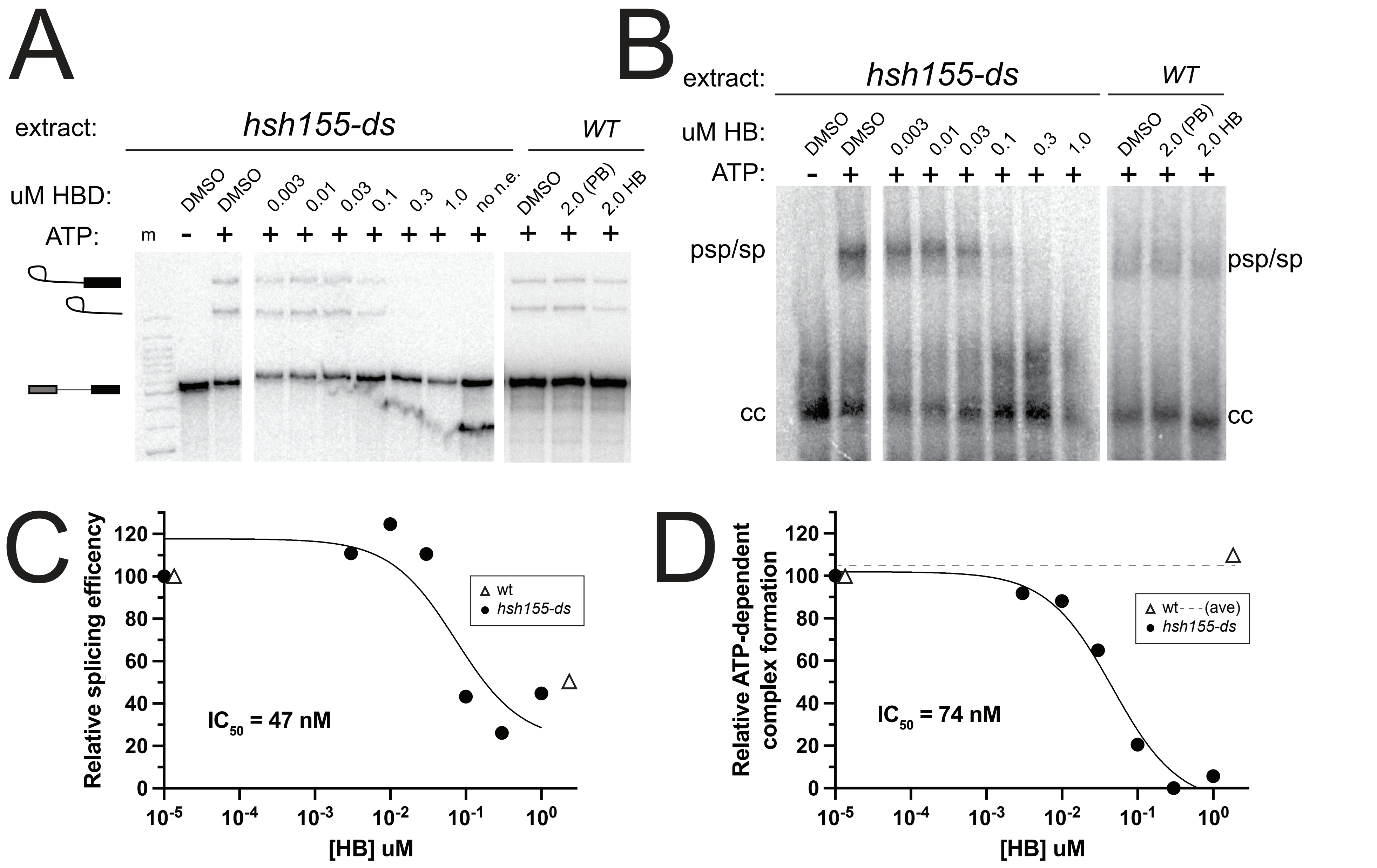


Hunter et al. (Ares) Fig S2

### Hunter_etal_FigS2E.pdf

Hunter et al. (Ares) Fig S2E

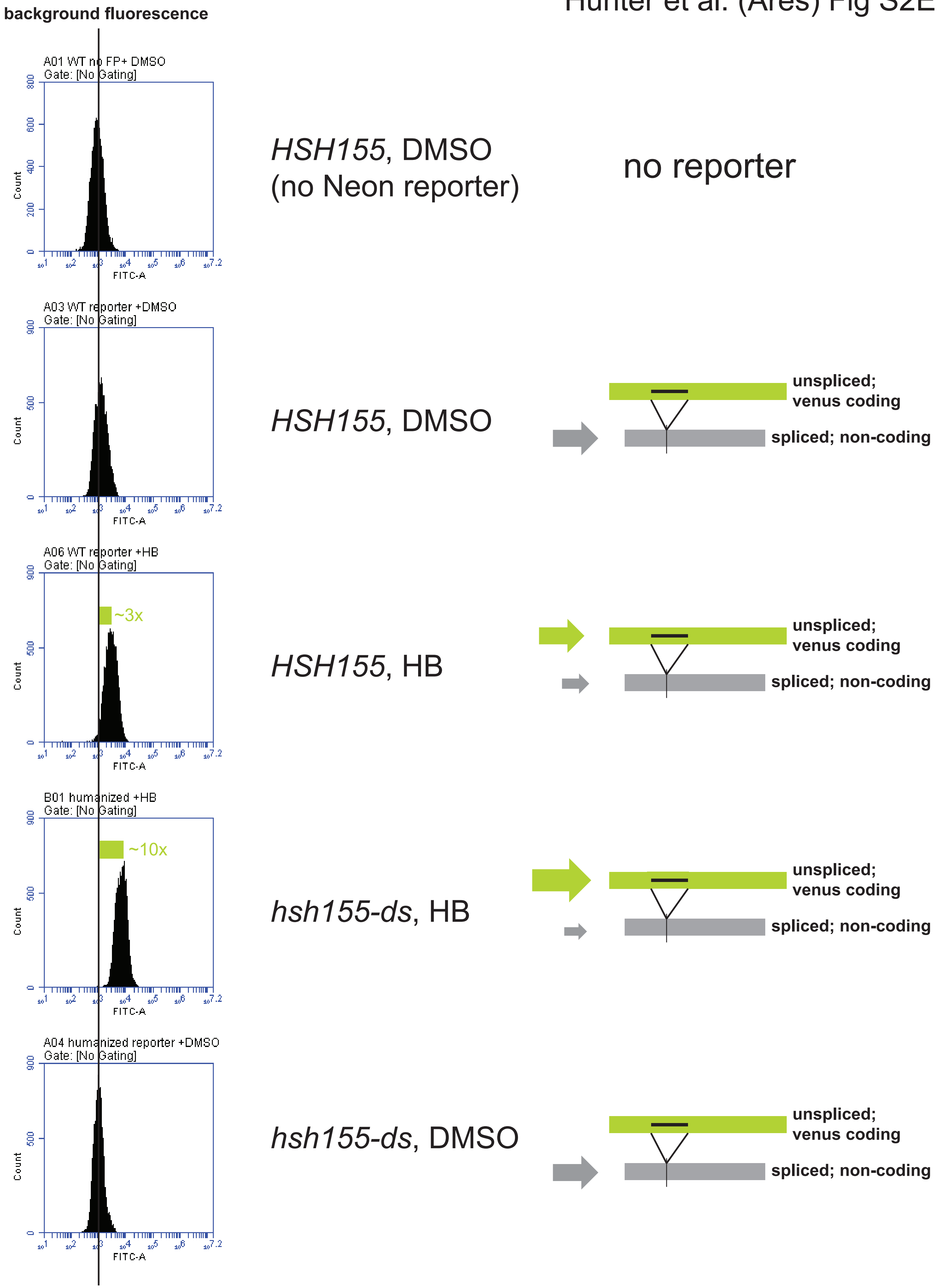

### Hunter_etal_FigS3A-F.pdf

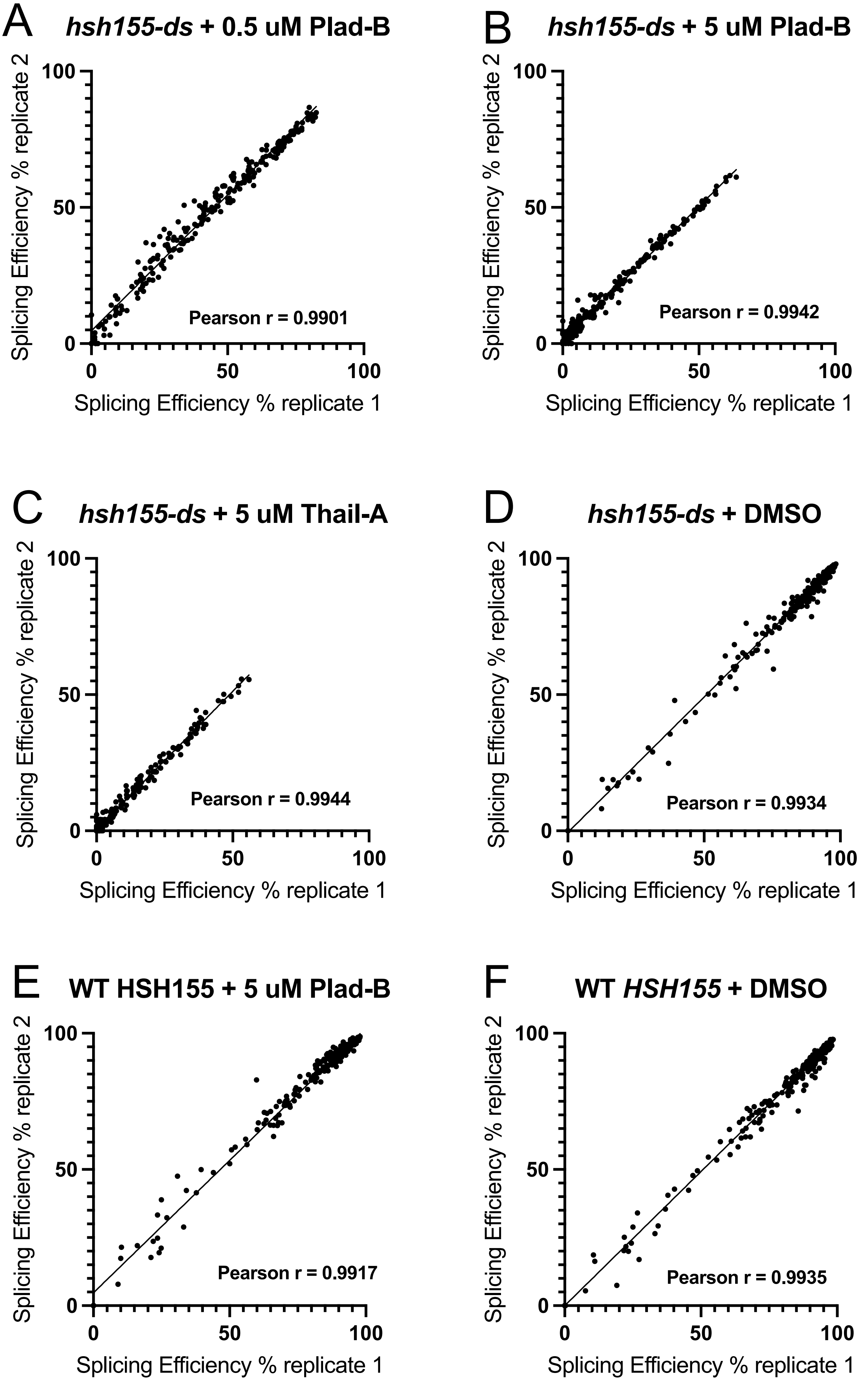

### Hunter_etal_FigS4.pdf

Hunter et al. (Ares) Fig S4

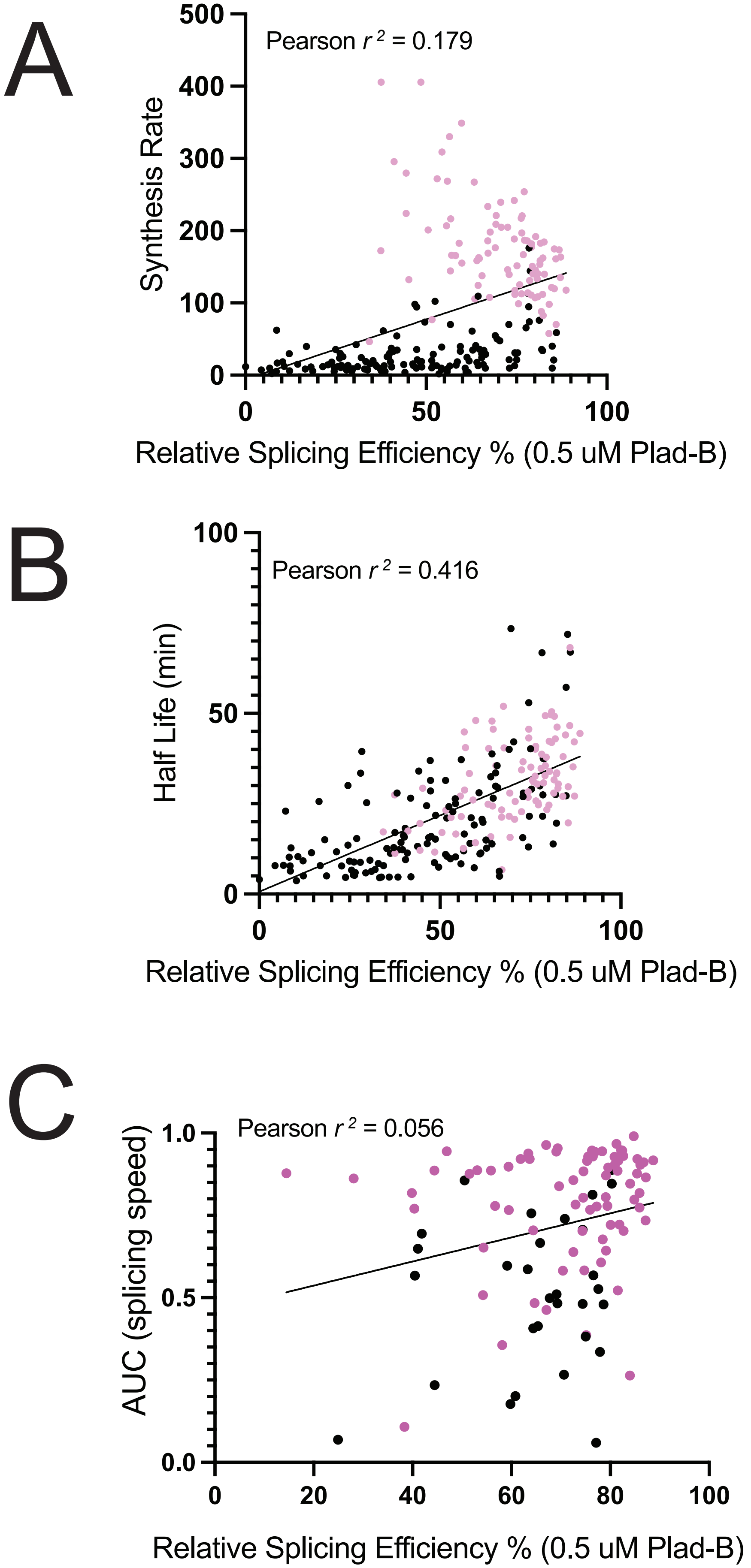
